# A new transgene mouse model using an extravesicular EGFP tag to elucidate the in vivo function of extracellular vesicles

**DOI:** 10.1101/2021.07.05.451120

**Authors:** Mikkel Ø. Nørgård, Lasse B. Steffensen, Didde R. Hansen, Ernst-Martin Füchtbauer, Morten B. Engelund, Henrik Dimke, Boye L. Jensen, Ditte C. Andersen, Per Svenningsen

**Author notes:** Correspondence Per Svenningsen, Dept. of Molecular Medicine, Cardiovascular and Renal Research, University of Southern Denmark, J.B. Winsloews vej 21.3, DK-5000 Odense C.

## Abstract

The *in vivo* function of cell-derived extracellular vesicles (EVs) is challenging to establish since cell-specific EVs are difficult to isolate. We therefore created an EV reporter using CD9 to display enhanced green fluorescent protein (EGFP) on the EV surface. CD9-EGFP expression in cells did not affect EV size and concentration, but allowed for co-precipitation of EV markers TSG101 and ALIX from cell-conditioned medium by anti-GFP immunoprecipitation. We created a transgenic mouse where CD9-EGFP was inserted in the inverse orientation and double-floxed, ensuring Cre recombinase-dependent EV reporter expression. We crossed the EV reporter mice with mice expressing Cre ubiquitously (CMV-Cre), in cardiomyocytes (AMHC-Cre) and kidney epithelium (Pax8-Cre), respectively. The mice showed tissue-specific EGFP expression, and plasma and urine samples were used to immunoprecipitate EVs. CD9-EGFP EVs was detected in plasma samples from CMV-Cre/CD9-EGFP and AMHC-Cre/CD9-EGFP mice, but not in PAX8-Cre/CD9-EGFP mice. On the other hand, CD9-EGFP EVs were detected in urine samples from CMV-Cre/CD9-EGFP and PAX8-Cre/CD9-EGFP mice, but not AMHC-Cre/CD9-EGFP, indicating that plasma EVs are not filtered to the urine. In conclusion, our EV reporter mouse model enables Cre-dependent EV labeling, providing a new approach to study cell-specific EVs in vivo and gain new insight into their physiological and pathophysiological function.

## Introduction

Extracellular vesicles (EVs) are nanosized membrane-bound vesicles that may function as mediators of cell-cell communication by transfer of cellular proteins, lipids, and nucleic acids^1^. The presence of EVs in biological fluids, such as urine and plasma, has also gained clinical interest since it facilitates the monitoring of physiological and pathophysiological processes using a minimal invasive technique. While EVs potentially enable the non-invasive interrogation of cells buried deep within an organism, the cell-specific isolation and tracking of EVs in vivo remains a challenge in physiological relevant settings, since EVs are secreted from almost every cells type^2, 3, 4, 5, 6, 7, 8, 9^. This hampers our ability to investigate the role of EVs in a physiological relevant context.

EVs are heterogeneous group of cell-released vesicles and the two main forms are exosomes, and microvesicles. Their biogenesis differs: microvesicles bud off directly from the plasma membrane, and exosomes are of endosomal origin and accumulates in intraluminal vesicles (ILVs) in multivesicular bodies (MVBs). EVs are, however, enriched in tetraspanins, such as CD63 and CD9^10, 11, 12^, which are transmembrane proteins widely distributed in the plasma membrane^13^, and considered valid markers of EVs^14^.

The tetraspanins have been widely used to create genetic strategies to label EVs through fusion of their N-and C-terminals^15, 16, 17, 18, 19^, to fluorescence reporter proteins or luciferase. These reporters enable tissue-specific signal exclusively from EVs, and thereby overcome major limitations of previous approaches using radioisotopes, fluorescent dyes and magnetic conjugated nanoparticles to lipophilic reagents to label EVs^7, 8, 9^. The lipophilic reagents can be released from the EVs and cause distribution of non-EV-associated fluorescent signal^12, 13^. Furthermore, in vivo injections of labeled EVs from in vitro cultured cells may not be representative for in vivo functions of EVs both regarding their often supra-physiological concentrations and since they only are representative from one single cell type. While N and C terminal fusion of reporter proteins to the tetraspanins enables tracking of cell-specific EVs, the intravesicular localization of the reporter proteins do not allow for affinity isolation of the labeled EVs from biological fluids in that the tetraspanin terminals are located inside Evs^13, 12^.

However, tetraspanins can be used to display fluorescent proteins on the EV surface^17^ and we hypothesized that fusion of EGFP to the C-terminal of a truncated form of the tetraspanin CD9, devoid of the large extracellular domain and the last transmembrane domain, would create fluorescence labeled EVs with affinity tags. By flanking the inverted EV reporter with loxP sites^20^, this would create a genetic switch that enabled Cre recombinase-dependent EV reporter expression in vivo, and allow both easy and reliable tracking and isolation of cell-specific EVs, enabling us to achieve new knowledge regarding the physiological and pathophysiological functions of EVs.

## Results

### Fusion of EGFP to CD9 enable affinity isolation of EVs

We designed an EV reporter protein by fusion of mouse CD9 truncated after the first 117 amino acids (the third transmembrane domain) to EGFP (Figure1A). The predicted molecular weight of the fusion protein is 41.5 kDa. Stable transfected epithelial M1 cells – termed M1-CD9-EGFP cells showed green fluorescent intracellular vesicles by fluorescence microscopy (Fig 1B), and western blotting showed EGFP expressing only in transfected cells (Figure 1C, whole blot figure S3). We noted that without addition of protease inhibitors to the lysis buffer EGFP was present as two bands: one at ∼37 kDa, slightly below the expected molecular weight of the fusion protein, and one at ∼27 kDa (Figure S3); however, cells and conditioned medium with addition of protease inhibitors showed the 37 kDa CD9-EGFP band (Figure 1C). Cell conditioned medium from M1 and CD9-EGFP cells has similar extracellular particle concentration and size distribution (Figure 1D), and PEG-precipitated conditioned medium contained EV markers ALIX and Flotilin-1, but no actin (Figure 1C, whole blots figure S4-S6). The CD9-EGFP was only detected in conditioned medium from CD9-EGFP cells. Furthermore, after immunoprecipitation western blotting showed EGFP expression in conditioned medium from cells containing our CD9-EGFP and associated EV marker TSG101 and ALIX after GFP immunoprecipitation (Figure 1E, whole blot figure S7-S9). Thus, EVs from CD9-EGFP expression cells can be isolated with GFP immunoprecipitation.

**Figure 1:**
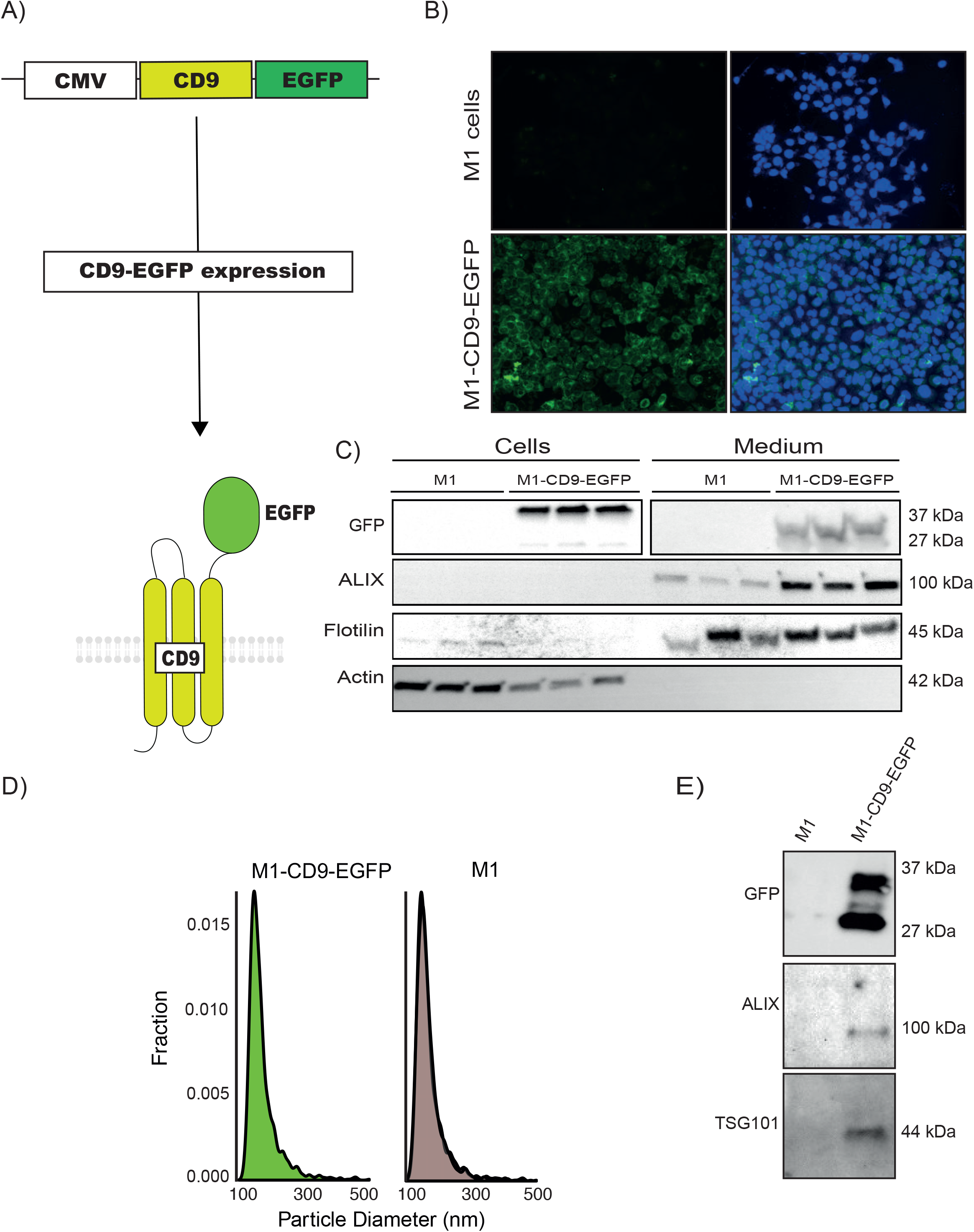
Expressing of EGFP and associated EV markers M1-CD9-EGFP cells and their conditioned medium. **(A)** Illustration of our EV reporter gene and protein. The CD9-EGFP coding sequence is driven by the CMV promoter and encodes a fusion protein consisting of the first 3 transmembrane domains of CD9 and EGFP, enabling genetic labeling and EV surface display of EGFP. (**B**) M1 cells stable transfected with CD9-EGFP are green fluorescent in contrast to non-transfected M1 cells. Nuclei (blue) and GFP (green). 20x magnification **(C)** M1 cells stable transfected with CD9-EGFP express a band reacting with an anti-GFP antibody in cells and conditioned medium. Actin was detected in cell lysates, and EVs markers ALIX and Flotillin in conditioned medium from transfected and non-transfected M1 cells. **(D)** Tunable resistive pulse sensing on conditioned medium from non-transfected M1 cells and stable transfected CD9-EGFP cells indicated that the size distribution of EVs is not affected the expression reporter proteins (n=3). **(E)** Anti-GFP immunoprecipitation of cells conditioned medium co-isolates CD9-EGFP and EV markers ALIX and TSG101 only in M1 cells stable transfected with CD9-EGFP (n = 3).

### Cre recombinase-dependent CD9-EGFP expression

To create cell-specific CD9-EGFP expression, we inverted the coding sequence of EV reporter protein CD9-EGFP and flanked it by double lox sites, allowing for CD9-EGFP expression driven by a CAG promoter only in Cre recombinase expressing cells (Figure 2A). HEK293T cells transient transfected with CD9-EGFP transient did not express CD9-EGFP; however, co-transfection with Cre recombinase yielded green fluorescent cells (Figure 2B) and CD9-EGFP expression in cells and condition medium (Figure 2C, whole blot figure S10-11). We created a transgene mouse through pro-nuclear injection of the linearized construct and obtained founder mice. We selected a founder mouse which showed expression of EGFP in Cre-expressing cardiomyocytes when crossed with AMHC-Cre mouse after tamoxifen injection (supplement figure 1A). After backcrossing onto a C59Bl/6 background, we isolated genomic DNA from the liver of a CD9-EGFP mouse and nanopore sequencing indicated insertion in the mouse genome at approximately chromosomal location 4:99.620.973. The insert was present in two copies in immediately adjacent to each other in opposite directions. The insert does not disrupt any known mouse genes at the location where it has been embedded. The area of insertion is, however, an annotated constrained conserved region between multiple eutherian mammals. We designed primers for genotyping (supplement figure 1B) that produced PCR products at 449 bp in wildtype mice and 324 bp in homozygotes (Figure 2D) enabling us to identify transgenic mice.

**Figure 2:**
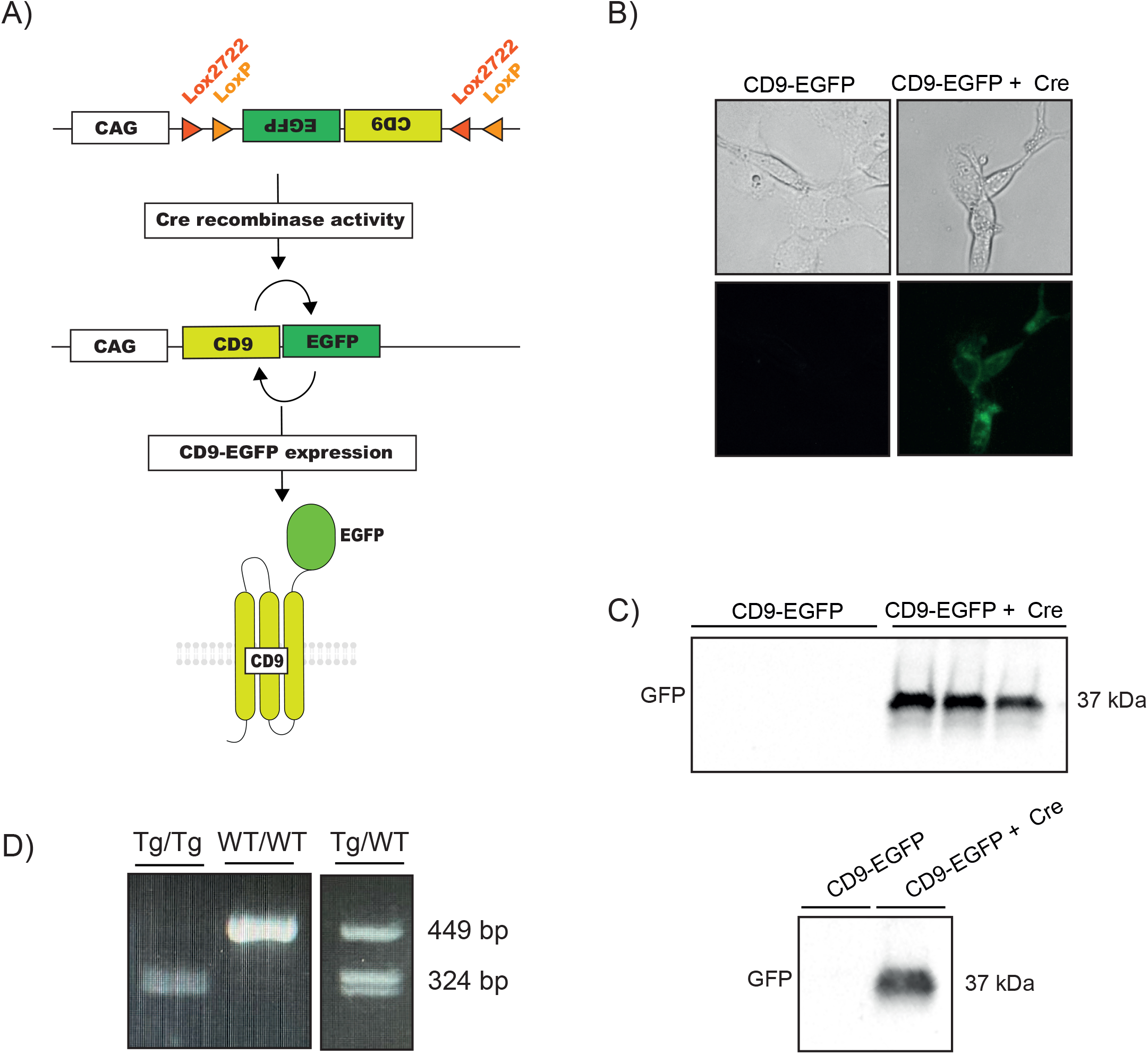
Cre recombinase-dependent expression of CD9-EGFP. **(A)** Our EV reporter protein CD9-EGFP was inverted and flanked by double lox sites. Upon Cre recombinase expression, the CD9-EGFP is inverted and yields CD9-EGFP expression driven by the CAG promoter. **(B)** HEK293T cells transiently transfected with double-floxed and inverted CD9-EGFP only express EGFP when co-transfected with Cre recombinase. **(C)** HEK293T cells co-transfected with double-floxed and inverted CD9-EGFP expressed CD9-EGFP in cells and conditioned medium. **(D)** The double-floxed and inverted CD9-EGFP construct is inserted into chromosome 4 of the EVRep mouse, and mice homo- (Tg/Tg) and heterozygous (Tg/WT) can be identified by PCR.

### Tissue-specific EGFP signal in different Cre-positive transgenic mice

We used established transgenic Cre recombinase mice to enable tissue-specific activation of the EV reporter protein by crossbreading with our EV Reporter (EVRep) mice. In CMV-Cre/CD9-EGFP mice, our mice crossed with CMV-mice expression Cre recombinase under the control of ubiquitiously active human cytomegalovirus (CMV) promoter, Cre positive mice showed EGFP expression in both kidney, liver, lung spleen, and heart by anti-GFP immunohistochemistry (IHC) (Figure 3A). In CMV-Cre/CD9-EGFP Cre negative mice EGFP was undetectable in all investigated tissues. To show more cell-specific expression, we crossed our EVRep mice with AMHC-Cre mice transgenic mice expressing tamoxifen-induced Cre recombinase activity under the control of the alpha-MHC promotor, and Pax8-Cre mice expressing Cre recombinase specific in the renal epithelium, respectively. In AMHC-Cre/CD9-EGFP mice injected with tamoxifen, EGFP signal was detected in the heart of Cre positive mice by western blotting and direct fluorescence microcopy (Figure 4A and B, whole blot figure S12). EGFP was undetectable in the kidney, liver, lung, and spleen in Cre recombinase positive mice and the heart of Cre recombinase negative mice (figure 4A). In the Pax8-Cre/CD9-EGFP mice, the western blotting indicates prominent CD9-EGFP expression in the kidneys specifically in Cre positive mice (Figure 5A, whole blots figure S13); however, green fluorescence was not directly detectable in the kidney epithelium by fluorescence microscopy of frozen kidney section, and only scarce EGFP signal was associated with glomerulies (Figure 5B). The preparation of frozen sections involved exposure of the tissue to large osmotic gradient, and we therefore tested whether epithelial EGFP expression was present in paraffin-embedded formalin fixed kidneys. Using an anti-EGFP antibody, we readily detected epithelial EGFP expression (Figure 5C). Thus, our EVrep mouse allow for Cre-dependent expression of CD9-EGFP.

**Figure 3:**
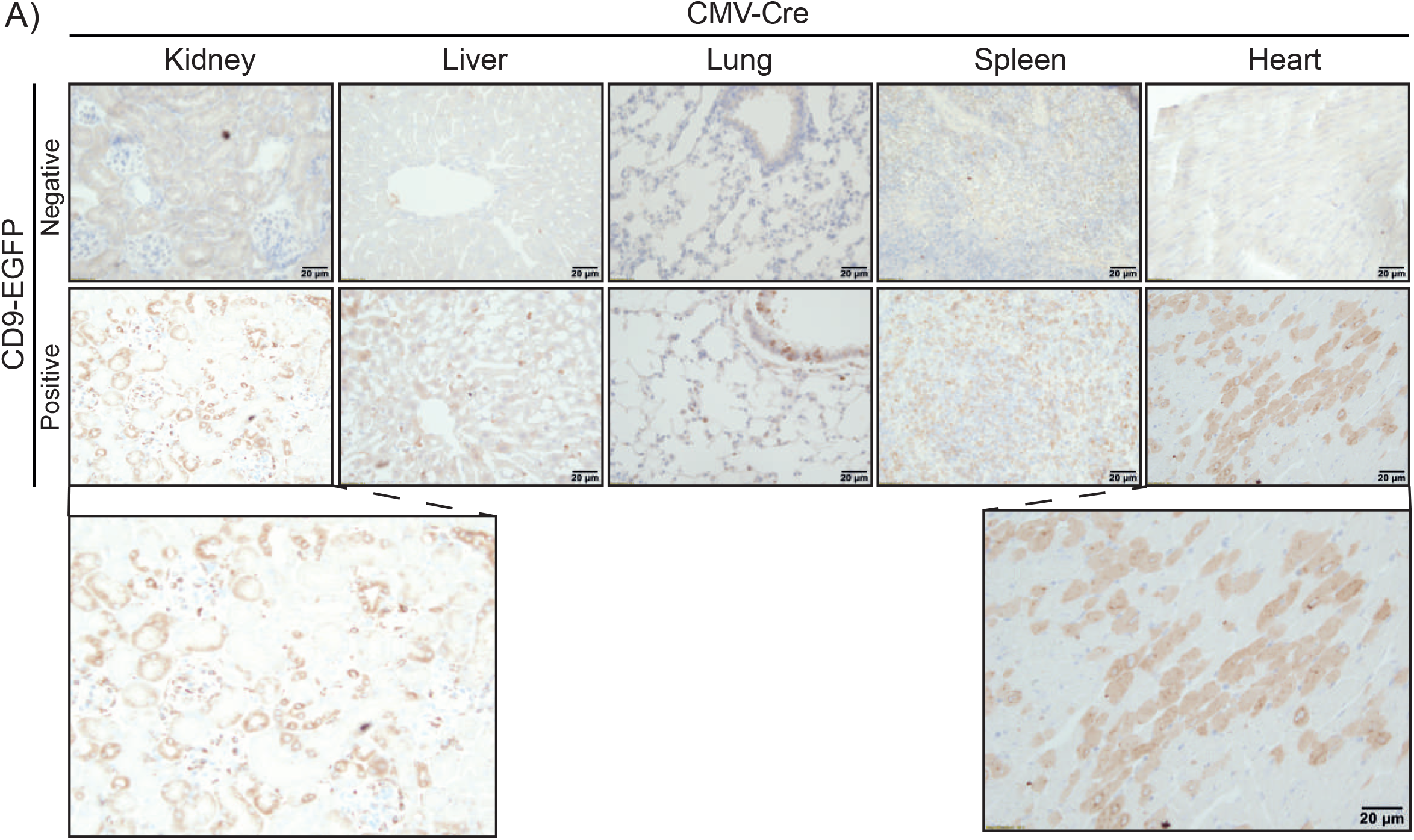
Expression of EGFP in tissue from CD9-EGFP positive CMV-Cre mice. **(A)** In contrast to CD9-EGFP negative CMV-Cre mice, EGFP expression is detected by anti-GFP (brown) immunohistochemical staining of paraffin-embedded kidney, liver, lung, spleen and heart in CD9-EGFP positive CMV-Cre mice.

**Figure 4:**
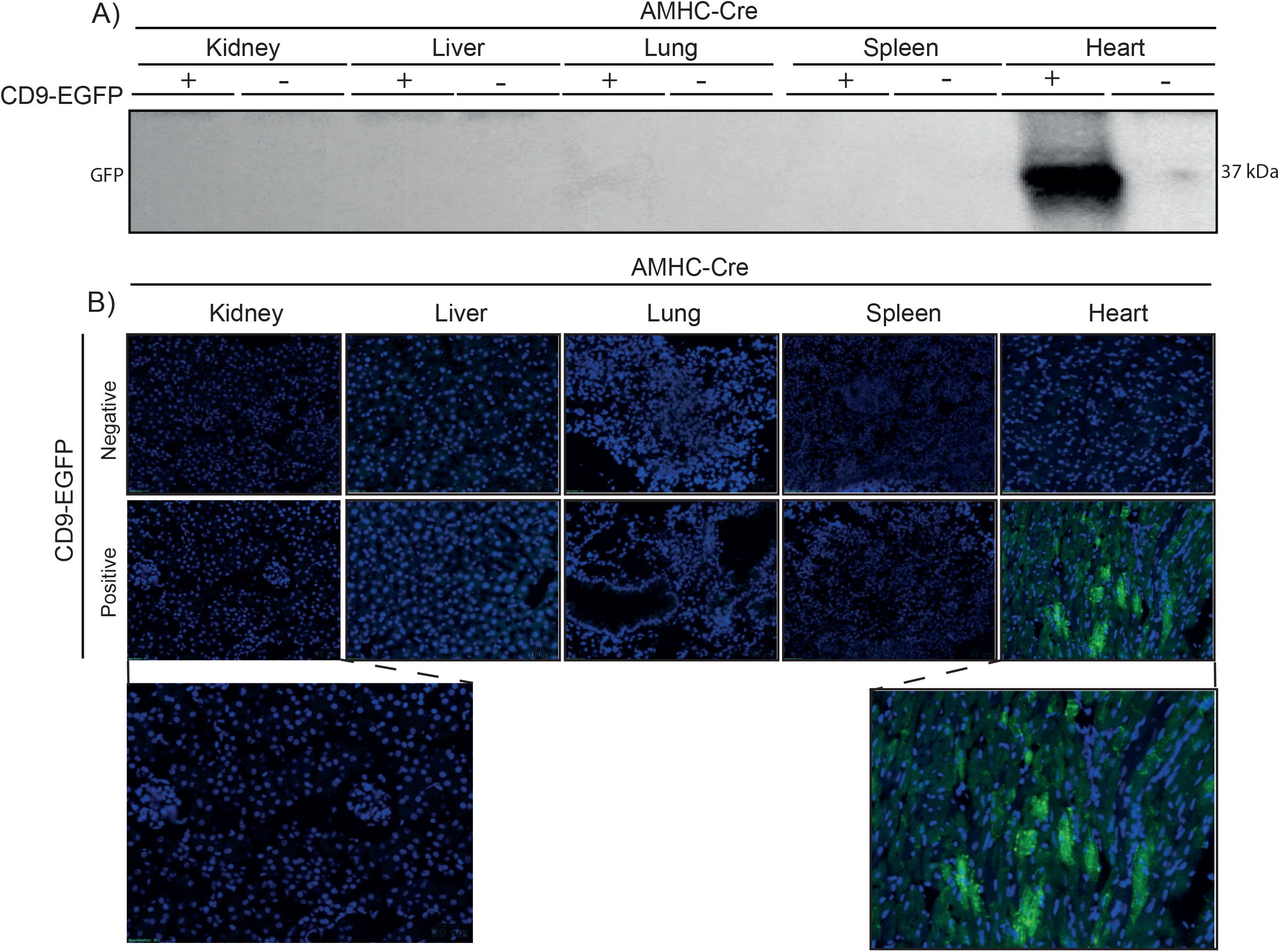
Expression of EGFP in cardiomyocytes from CD9-EGFP positive AMHC-Cre mice. **(A)** Tissue homogenates from AMHC-Cre mice display CD9-EGFP expression only in heart from CD9-EGFP positive mice. **(B)** Tissue sections from AMHC-Cre mice only shows EGFP expression (green) in cardiomyocytes in CD9-positive mice. Nuclei (blue).

**Figure 5:**
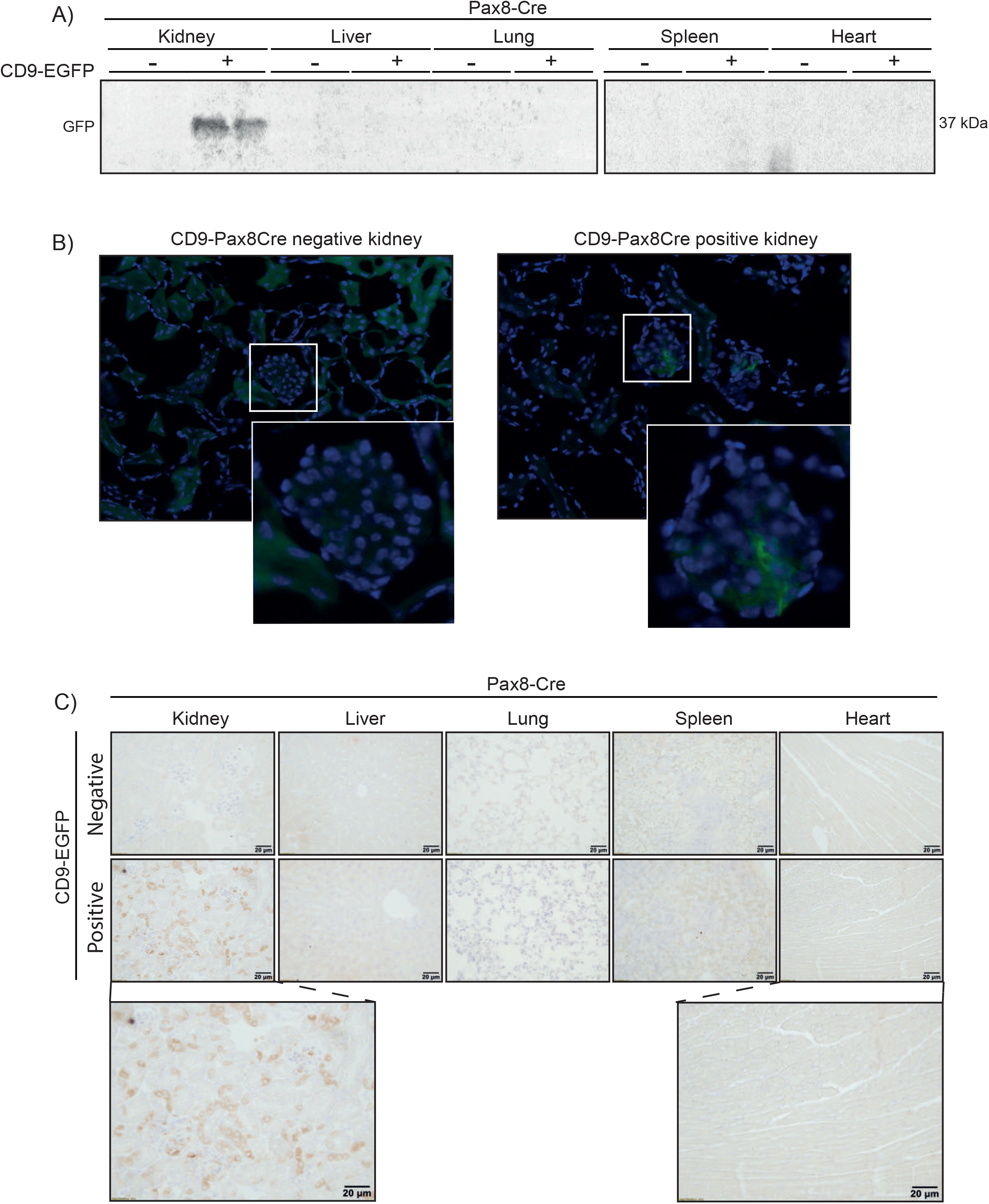
Expression of EGFP in kidney epithelial cells from CD9-EGFP positive Pax8-Cre mice. **(A)** Tissue homogenates form Pax8-Cre mice display CD9-EGFP expression in kidneys from CD9-EGFP positive mice. **(B)** Direct fluorescence imaging of kidneys from Pax8-Cre/CD9-EGFP mice only shows EGFP expression (green) in glomerular cells. Nuclei (blue). **(C)** Immunohistochemical staining of paraffin-embedded kidney, liver, lung, spleen and heart from Pax8-Cre mice, EGFP expression is detected by anti-GFP (brown) in epithelial cells in kidney from CD9-EGFP positive mice.

### Isolation of EGFP-containing extracellular vesicles from plasma and urine

EVs have been detected in most body fluids, and we used the different cell-specific EV mice to analyze for their contribution of EVs in plasma and urine samples. Plasma samples were PEG-precipitated and EGFP was immunoprecipitated using anti-GFP nanobody conjugated beads. CD9-EGFP was detected from CMV-Cre/CD9-EGFP and AMHC-Cre/CD9-EGFP positive mice Cre recombinase positive mice, while CD9-EGFP was undetected in Pax8-Cre/CD9-EGFP positive or CD9-EGFP negative transgenic mice (Figure 6A, whole blot figure S14). We tested the plasma samples from the PAX8-Cre and AMH-Cre mice for co-precipiation of EV markers, and found that ALIX co-precipitated with CD9-EGFP only in AMC-Cre/CD9-EGFP positive mice (Figure 6A, whole blot figure S15). In contrast to the plasma samples, urine samples from CMV-Cre and CD9-Pax8-Cre mice, but not AMHC-Cre/CD9-EGFP and CD9-EGFP negative mice, contained EGFP and CD9-EGFP (Figure 6B, whole blot figure S16). The affinity isolation of EGFP, co-precipitated EV marker CD81 in Pax8-Cre/CD9-EGFP postive mice, but not in AMHC-Cre/CD9-EGFP positive and CD9-EGFP negative mice (Figure 6B, whole blot figure S17).

**Figure 6:**
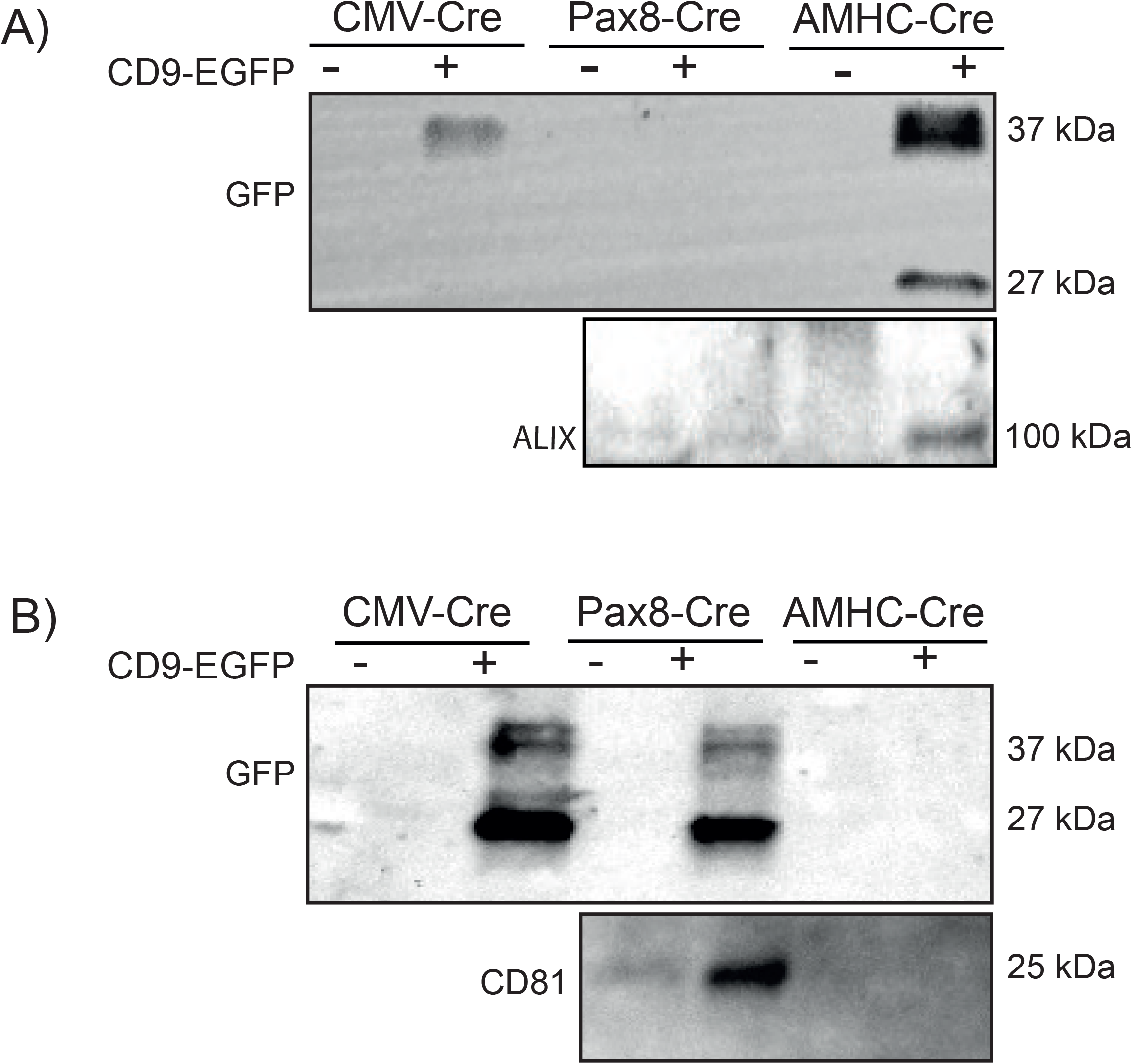
Isolation of EVs from plasma and urine. **(A)** Anti-GFP precipitation of plasma samples from CMV-Cre, Pax8-Cre and AMHC-Cre mice only isolated CD9-EGFP from CD9-EGFP positive CMV-Cre and AMHC-Cre mice. The CD9-EGFP precipitation co-isolated EV marker ALIX in plasma samples from CD9-EGFP positive AMHC-Cre mice. **(B)** CD9-EGFP is only detected by anti-GFP immunoprecipitation of urine samples from CD9-EGFP positive CMV-Cre and Pax8-Cre mice, but not AMHC-Cre. CD9-EGFP was co-precipitated with EV marker CD81 in Pax8-Cre/CD9-EGFP positive mice.

## Discussion

Using a genetically encoded CD9-EGFP fusion protein that display EGFP on EVs combined with a Cre-dependent switch, we have created a new epithelial cell line and transgenic mouse enabling easy isolation of cell-specific EVs from plasma and urine. This model is the first genetic EV reporter model that both allow easy tracking and isolation of EVs in vivo in mice and we believe this novel tool will allow us to gain new insight into EV function.

EVs isolated from biological fluid provide access to cell-derived lipids, RNA and proteins; however, the representation of EVs from different cell types in for example urine and plasma is not well described. Consistent with previous a study^22^, we found that EVs from cardiomyocytes was readily detectable in plasma samples. Intravascular injection of labelled EV have been detected in urine^21^; however, we did not detect cardiomyocyte-derived EVs in urine, indicating that EVs are not glomerular filtered in the kidney. One the other hand, we detected kidney epithelial EVs urine. This is consistent with our previous finding using proteomic database analysis showing that 99.96 % of urinary EV-associated proteins are likely to originate from kidney, the urinary tract epithelium and the male reproductive tract in humans^23^. Thus, the distribution of EVs appears to be restricted and EVs isolated from different fluid compartments, e.g., plasma versus urine, may only represent a subset of cell-types.

We found that kidney epithelial-derived EV signal was significantly affected by preparation of frozen kidney cross-sections before fluorescence microscopy, while cardiomyocyte EV tissue abundance was less affected. Western blotting suggested that CD9-EGFP was abundantly expressed in Pax8-Cre/CD9-EGFP mice, but in frozen kidney sections the EGFP was only slightly visible in glomerulus of Cre recombinase positive kidney by directed fluorescence microscopy. Although the reason for this discrepancy is not known. Kidney epithelial cells are, however, highly water permeable^24^, and we speculate that the osmotic changes, e.g., 25% sucrose, imposed during tissue during preparation are involved. Recently, it was shown that the handling and preparation of tissue samples significantly affect tissue EV abundance through release to e.g. washing buffers^25^. Our observations agree with this and suggest that careful and specific tissue preparation is important when analyzing tissue EV levels and intercellular communication.

Cell-to-cell EV communication has already been demonstrated in several settings. In vitro, the transport of miRNA in EVs act as a neuron-to-astrocyte communication pathway in the central nervous system^26^ and stem cell-derived EVs appear to target bone marrow and peripheral sites^27^. In vivo, intravascscular administration of mesenchymal stem cell-derived EVs (MSC-EV) labelled with PKH26 dye closely resembles the positive effect of MSCs on postischemic recovery after acute tubular injury^28^. Other examples include i.v. injection of endothelial progenitor cell-derived EVs labelled with PKH26 dye isolated from cell-conditioned medium that protects from complement-mediated mesangial injury after experimental anti-Thy1.1 glomerulonephritis^29^. All together this suggest that EVs acts as the functional part. However, these injections cause a supra-physiological concentration of EVs from a single cell type, and the naturally occurring in vivo concentrations might not be sufficient to cause similar effects. Furthermore, important paracrine and autocrine effects might be altered or unobserved with this approach and affect the in vivo distribution and the biological function of the EVs. Importantly, injected EVs often accumulate in the liver, spleen and lungs regardless of the origin^28, 29, 30^. We did not observe CD9-EGFP accumulation in these organs under normal physiological circumstances. These observations emphasize some of the hurdles with injections of labeled EVs^31, 32^, and the use of a mouse model with genetically labeled cell-specific EVs will enable a more faithful mapping of the EV cell-to-cell communication.

Similar to our approach, several other studies have used genetic labeling of EVs ^33, 15, 18^ to track the faith of endogenous EVs. The approaches share the limitations that there is a risk that only a subpopulation of EVs is labeled. The number of identified EV subtypes is continuously growing and there is no consensus of which protein markers that represent the different population. The use of the genetic labeling may therefore add a bias in the downstream analysis.

Nonetheless, the ease at which cell-specific EVs can be isolate is a clear benefit, and allow for a deeper characterization and identification of subtype and cell-specific EV markers.

In summary, our novel transgenic EVRep mouse allows easy in vivo tracking and isolation of EVs and can be used to elucidate the physiological and pathophysiological mechanisms and their role as mediators of cell-cell-signaling.

## Methods

### Plasmids

The genes encoding truncated CD9 fused to EGFP and the doubled-floxed and inverted version was synthesized and cloned into plasmid pcDNA3.1 by BioCat GMBH. The coding sequence is shown in Supplement Figure 2A. The double-floxed, inverted CD9-EGFP from the pcDNA3.1 plasmid was amplified by PCR and ligated into pCAG-Cre (a gift from Connie Cepko; Addgene plasmid # 13775) digested with EcoRI and NotI (New England Biolabs). All constructs were verified by Sanger sequencing at Eurofins Genomics.

### Cell cultures

Mouse epithelial M1 cells were cultured as previously described^34^. For analysis of cell conditioned medium, PC1 serum-free medium (Lonza, No. 344018, Walkersville, MD, USA) was used. The M1 cells with stable CD9-EGFP expression was generated by transfection with pcDNA3.1 CD9-EGFP using Metafectene Pro (Biontex) and selection with G418 (InvivoGen). HEK293T cells were maintained in Dulbecco’s Modified Eagle Medium, F-12 Nutrient Mixture (Gibco, Sigma Aldrich, Denmark) containing 10% fetal bovine serum (FBS) and 1% Penicillin-Streptomycin in a 5% CO2 humidified incubator at 37° C.

### Animal Models

We linearized pCAG-DIO-CD9-EGFP with HindIII and SalI (New Engaldn Biolabs) and gel extracted (New England Biolabs) the 3649-bp fragment. The purified fragment was used to generate transgenic mice with genomic integration of the CAG-promoter driven double-floxed, inverted CD9-EGFP by pro-nucleus injection^35^. The animal experiment was performed in accordance with Danish Law under animal experimental permits #2019-15-0201-01644 and 2019-15-0202-00052. (Description of AMHC, CMV, Pax8-Cre).

### Sample collection

7-9 weeks old mice were transferred individually to metabolic cages (12:12h light-dark cycle, 28±1 °C) for three days with free access to water and normal rodent diet (LabDiet® 5001, Forth Worth, TX, USA). After 24h acclimation urine samples were collected on day two and three. Subsequently, mice were used for perfusion fixation or direct organ harvest as described below. Urine was stored at -80 °C and a protease inhibitor in the ratio 1:1000 (P8340, Protease Inhibitor Cocktail, Sigma Aldrich, Denmark) was added when thawed. Mice used for immunohistochemistry and fluorescent microscopy were anesthetized by i.p. injection with 10 mg/kg Xylazine (Rompun, Bayer Healthcare, Shawnee Mission, KS) and 50 mg/kg Ketamine (Ketalar, Pfizer, Sandwich, Kent, UK). Blood samples were taken through the apex, before mice were flushed with PBS and fixed with 4% paraformaldehyde by retrograde perfusion via the left ventricle. Subsequently, mice were fixed for an additional 6 hours in paraformaldehyde and transferred to 1x PBS. Mice used for Western blot were likewise anesthetized, blood samples were taken and relevant organs were removed and snap-frozen in liquid nitrogen.

### Immunohistochemistry and fluorescence microscopy on cross-sections

Staining was performed on 2 μm paraffin-embedded tissue samples deparaffined in xylene and rehydrated in decreasing ethanol solutions (99%-70%) followed by target retrieval in heated TEG-buffer (1 mmol/L Tris, 0.5 mM EGTA, pH 9.0). After cooling, slides were exposed to a 50 mM NH_4_Cl and 0.3% H_2_O_2_ solution for 10 minutes, washed in 1x PBS, and incubated in 1x PBS with 0.3% Triton for 30 minutes at room temperature. Next, the slides were incubated with primary antibody (Anti-GFP) diluted in 1x PBS with 0.3% Triton x100 at 4°C overnight.

Next, slides were incubated for 30 minutes at room temperature, washed in 1x PBS + 0.05% tween five times and incubated with secondary HRP antibody in PBS + 0.05% tween for 1 h at room temperature. Slides were then washed in PBS and HRP was visualized by DAB-staining (3,3’-diaminobenzidine). Lastly, slides were stained in hematoxylin mounted with Aquatex (Merck KGaA, Darmstadt, Germany).

For fluorescence microscopy, snap-frozen organs were removed from the mice placed in 1x PBS with 0.05% acid and 25% sucrose overnight. Next, organs were placed in 1x PBS with 0.05% acid and 50% sucrose for 2 hours, embedded in OCT tissue freezing medium and 5 μm cross-sectioned on a Cryostat (Leica CM3050 S, Leica Biosystems, USA). Cross-sections were placed in 1x TBS for 10 minutes, dried and one drop of Slowfade^™^ Gold antifade reagent with DAPI (Invitrogen, Thermo Fisher Scientific, Eugene, OR USA) was added.

### Isolation of extracellular vesicles from cells and medium

48h before harvest, cell medium was changed PC1 serum-free medium (Lonza, No. 344018, Walkersville, MD, USA). Cells were lysed in RIPA Lysis buffer for 1h at 4° C on an orbital shaker. Next, 25 cm cell scrapers were used to release cells from the flask into the RIPA buffer. The solution was then centrifugated at 13000 G at 4° C for 10 min and the supernatant was stored in new tubes on -80° C.

Medium was transferred to 15 mL tubes and centrifugated 10 min at 5000 G. The supernatant was then transferred to new tubes and stored at -80° C. To isolate extracellular vesicles, thawed medium was centrifugated for 15 min at 5000 G at 4° C. The supernatant was mixed with an equal amount of freshly made ExtraPEG (16% PEG-6000, Sigma Aldrich, 1 M NaCl, Milli-Q water) and left in a Multi-Rotator at 4° C overnight. Next, samples were centrifugated for 15 min at 5000 G at 4° C, and the supernatant was discarded. The remaining pellet was resuspended in 100 μl x1 RIPA buffer and stored at -20° C until further use.

The use of a Polyethylene Glycol-based method (ExtraPEG) for enrichment of EVs is rapid and inexpensive based on precipitation of EVs and other compounds in the solution of super hydrophilic polymers, PEGs.^36, 37^. The amount and quality of isolated EVs are comparable to commercial reagents^38^.

### Immunohistochemistry on cells

Cells were seeded on coverslips in a multiple 12-wells plate (Biocoat Cell environments, Poly-D-Lysine Cellware) and given 24 h to attach to the coverslips. Next, cells were fixed in 4% Paraformaldehyde for 10 min and rinsed 2 times in PBS with 1 mM MgCl_2_ and 0.1 mM CaCl_2_, followed by permeabilization with 0.3% triton in PBS for 15 min and another wash in PBS with 1 mM MgCl_2_ and 0.1 mM CaCl_2_.

Afterwards, cells were incubated with 4’,6-diamidino-2-phenylindole (D9542-10MG DAPI, Sigma Aldrich, Denmark) to stain DNA followed by 5 times wash in PBS. Lastly, coverslips containing cells were mounted with fluorescent mounting media (DAKO, Carpinteria, CA, USA) on coverslips. ImageJ, version 2.0.0-RC-43/1.10e was used to analyze pictures.

### Immunoblotting

Samples were mixed with LDS sample buffer (NuPAGE, Invitrogen, Thermo Fischer Scientific, Van Allen Way, Carlsbad, USA) and sample reducing agent (NuPAGE, Invitrogen, Thermo Fischer Scientific, Van Allen Way, Carlsbad, USA) and heated for 10 min before they were ran on a gel. The gel was then transferred to a membrane activated in 99% Ethanol. Afterwards, the membrane was blocked for 30 min in 5% skimmed milk and incubated with primary antibody overnight at 4° C (supplement 1C). Following, membranes were washed 3 times in Tris-buffered saline with tween-20 (TBST, 20 mM Tris-base, 137 nM NaCl, 0.05 % Tween-20 (Merck), pH=7.6) and incubated with secondary antibody (supplement 1C) for 1h at room temperature. Next, membranes were washed 3 times TBST, and proteins were visualized by ECL plus a Molecular Imager (ChemiDoc XRS+, BIO-RAD) with Image Lab software (BIO-RAD).

### Immunoprecipitation of EGFP associated EVs

EVs containing EGFP were precipitated from 200 μl plasma and 750 μl urine with extraPEG as described above, except that the pellet was resuspended in 100 μl PBS to keep EVs intact. EVs were further up-concentrated with ChromoTek GFP-trap^®^ Magnetic Beads (Chromotek GmbH, Germany). 25 μl beads were added to a 1.5 ml tube and rinsed with 1000 μl ice-cold PBS. Next, plasma or urine were added to the equilibrated beads and tubes were placed in a Multi-Rotator (Grant-bio, PTR-35) at 4° C for 1 h. Samples were when placed back in the magnet and allowed to attach before the supernatant was discarded. Subsequently, beads were resuspended in 1 mL ice-cold PBS-T and the samples were placed back in the magnet. The supernatant was discarded after 3 min when the beads were attached to the magnet, and this step was repeated 4 times.

### Tunable Resistive Pulse Sensing

A qNano platform with a Nanopore NP150 (Izon Science, Oxford, UK) and polystyrene calibration beads CPC200 (Izon Science), Oxford, UK) were used for calibration of relative particle size and speed. 35 μl cell media was loaded, and analyses were performed according to manufactory instructions.

### Nanopore sequencing

Mouse liver from a CD9-EGFP positive mouse was homogenized and DNA was extracted using the Nanobind Tissue Big DNA Kit (Circulomics, Baltimore, MD, USA) according to the manufacturer’s instructions. The DNA was prepared for Oxford Nanopore sequencing using the Ligation Sequencing Kit (Oxford Nanopore Technologies, Oxford, UK). The prepared library was then sequenced using one flow cell on the Oxford Nanopore PromethION sequencing platform. Base-calling was performed with MinKNOW software and FASTQ files were aligned to a custom mouse genome containing the CD9EGFP insert with various promoter sequences and accessory sequences as an extra chromosome.

Alignment was performed using Minimap2 (Li, 2018). Data were visualized using IGV ver2.8.1 (Broad Institute), Genome Ribbon (Nattestad et al., 2020), and NCBI BLAST.

## Notes

### Competing Interest Statement

The authors have declared no competing interest.

