## Supplement for "A new transgene mouse model using an extravesicular EGFP tag to elucidate the in vivo function of extracellular vesicles"

A)

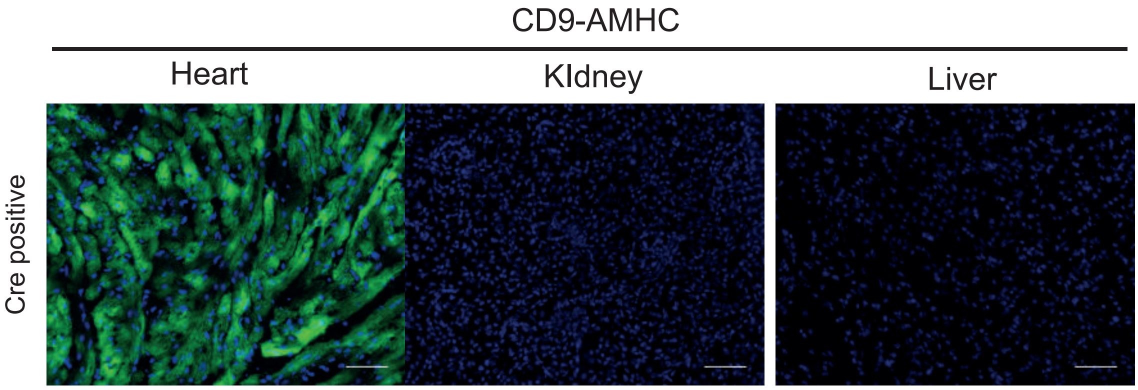

B)

Table 1: Primer sequences used for genotyping

| ID | Sequence | Annealing temperature C° | Function | BP |
|---|---|---|---|---|
| PS0819 | TGCACCTCAA ACACTCAAGC | 60 | Genotypning af CD9-EGFP forward | 449/324 |
| PS0820 | TCTTCCTGGAA CACAGCTCA | 60 | Genotypning af CD9-EGFP reverse 449 bp product in wildtype and hetero | 449 |
| PS0821 | AGCCAGATTTT TCCTCCTCTC | 60 | Genotypning af CD9-EGFP 324 bp product in CD9 and hetero (with PS0819) | 324 |

C)

Table 2: Primary and sencondary antibodies

| Target protein | Company | Catalog # | Dilution |
|---|---|---|---|
| **Western blot** | **Primary Antibodies** | | |
| **GFP** | Nordic BioSite, SE | GTX113617 | 1:1000 |
| **ALIX** | Cell Signaling Technology, NL | 2171 | 1:1000 |
| **Flotilin** | Abcam, UK | Ab13493 | 1:1000 |
| **Calnexin** | ECM Biosciences, USA | CMA4371 | 1:1000 |
| **Beta-actin** | Abcam, UK | Ab8227 | 1:1000 |
| **CD81** | Santa Cruz Biotechnology, USA | E1817 | 1:1000 |
| **Immunohistochemistry** | **Primary Antibodies** | | |
| **Anti-GFP** | Abcam, UK | Ab6673 | 1:200 |
| **Secondary Antibodies** | | | |
| **Polyclonal Goat anti-mouse immunoglobin/HRP** | | DAKO, Denmark | P0447 |
| **Polyclonal Goat anti-rabbit immunoglobin/HRP** | | DAKO, Denmark | P0448 |
| **Polyclonal Rabbit Anti-Goat immunoglobulins/HRP** | | DAKO, Denmark | P0449 |

A)

CD9-EGFP-Flag

atgccggtcaaaggaggtagcaagtgcatcaaatacctgctcttcggatttaacttcatcttctggctcgctggcattgcagtgcttgctattggactatggct
ccgattcgactctcagaccaagagcatcttcgagcaagagaataaccattccagtttctacacaggagtgtacattctgattggagccggggccctcatgat
gctggttggtttcctgggctgctgtggagctgtacaagagtcccagtgcatgctgggattgttcttcgggttcctcttggtgatattcgccattgagatagccg
ccgccgtctggggctatacccacaaggatgaggtgattaaaatggtgagcaagggcgaggagctgttcaccggggtggtgcccatcctggtcgagctgga
cggcgacgtaaacggccacaagttcagcgtgtctggcgagggcgagggcgatgccacctacggcaagctgaccctgaagttcatctgcaccaccggcaa
gctgcccgtgccctggcccaccctcgtgaccaccctgacctacggcgtgcagtgcttcagccgctaccccgaccacatgaagcagcacgacttcttcaagtc
cgccatgcccgaaggctacgtccaggagcgcaccatcttcttcaaggacgacggcaactacaagacccgcgccgaggtgaagttcgagggcgacaccct
ggtgaaccgcatcgagctgaagggcatcgacttcaaggaggacggcaacatcctggggcacaagctggagtacaactacaacagccacaacgtctatat
catggccgacaagcagaagaacggcatcaaggcgaacttcaagatccgccacaacatcgaggacggcagcgtgcagctcgccgaccactaccagcaga
acacccccatcggcgacggccccgtgctgctgcccgacaaccactacctgagcacccagtccgccctgagcaaagaccccaacgagaagcgcgatcaca
tggtcctgctggagttcgtgaccgccgccgggatcactctcggcatggacgagctgtacaaggcagcaaatgatatcctggattacaaggatgacgacgat
aaggtttaa

CD9
EGFP
Flag tag
Stop codon

Double floxed inverted CD9-EGFP

ataacttcgtataggatactttatacgaagttatgcagaatggtagctggattgtagctgctattagcaatatgaaacctcttaataacttcgtatagcatac
attatacgaagttatggcgcgccctaTTAAACCTTATCGTCGTCATCCTTGTAATCCAGGATATCATTTGCTGCCTTGTACAGCTC
GTCCATGCCGAGAGTGATCCCGGCGGCGGTCACGAACTCCAGCAGGACCATGTGATCGCGCTTCTCGTTGGGGTCTTT
GCTCAGGGCGGACTGGGTGCTCAGGTAGTGGTTGTCGGGCAGCAGCACGGGGCCGTCGCCGATGGGGGTGTTCTGC
TGGTAGTGGTCGGCGAGCTGCACGCTGCCGTCCTCGATGTTGTGGCGGATCTTGAAGTTCGCCTTGATGCCGTTCTTC
TGCTTGTCGGCCATGATATAGACGTTGTGGCTGTTGTAGTTGTACTCCAGCTTGTGCCCCAGGATGTTGCCGTCCTCCT
TGAAGTCGATGCCCTTCAGCTCGATGCGGTTCACCAGGGTGTCGCCCTCGAACTTCACCTCGGCGCGGGTCTTGTAGT
TGCCGTCGTCCTTGAAGAAGATGGTGCGCTCCTGGACGTAGCCTTCGGGCATGGCGGACTTGAAGAAGTCGTGCTGC
TTCATGTGGTCGGGGTAGCGGCTGAAGCACTGCACGCCGTAGGTCAGGGTGGTCACGAGGGTGGGCCAGGGCACG
GGCAGCTTGCCGGTGGTGCAGATGAACTTCAGGGTCAGCTTGCCGTAGGTGGCATCGCCCTCGCCCTCGCCAGACAC
GCTGAACTTGTGGCCGTTTACGTCGCCGTCCAGCTCGACCAGGATGGGCACCACCCCGGTGAACAGCTCCTCGCCCTT
GCTCACCATTTTAATCACCTCATCCTTGTGGGTATAGCCCCAGACGGCGGCGGCTATCTCAATGGCGAATATCACCAAG
AGGAACCCGAAGAACAATCCCAGCATGCACTGGGACTCTTGTACAGCTCCACAGCAGCCCAGGAAACCAACCAGCAT
CATGAGGGCCCCGGCTCCAATCAGAATGTACACTCCTGTGTAGAAACTGGAATGGTTATTCTCTTGCTCGAAGATGCT
CTTGGTCTGAGAGTCGAATCGGAGCCATAGTCCAATAGCAAGCACTGCAATGCCAGCGAGCCAGAAGATGAAGTTAA
ATCCGAAGAGCAGGTATTTGATGCACTTGCTACCTCCTTTGACCGGCATggtggctagcataacttcgtataaagtatcctatacgaa
gttatttgccttaacccagaaattatcactgttattctttagaatggtgcaaagaataacttcgtataatgtatgctatacgaagttat

lox2722
loxP
Inverted CD9
Inverted EGFP
Inverted Flag tag
Stop codon

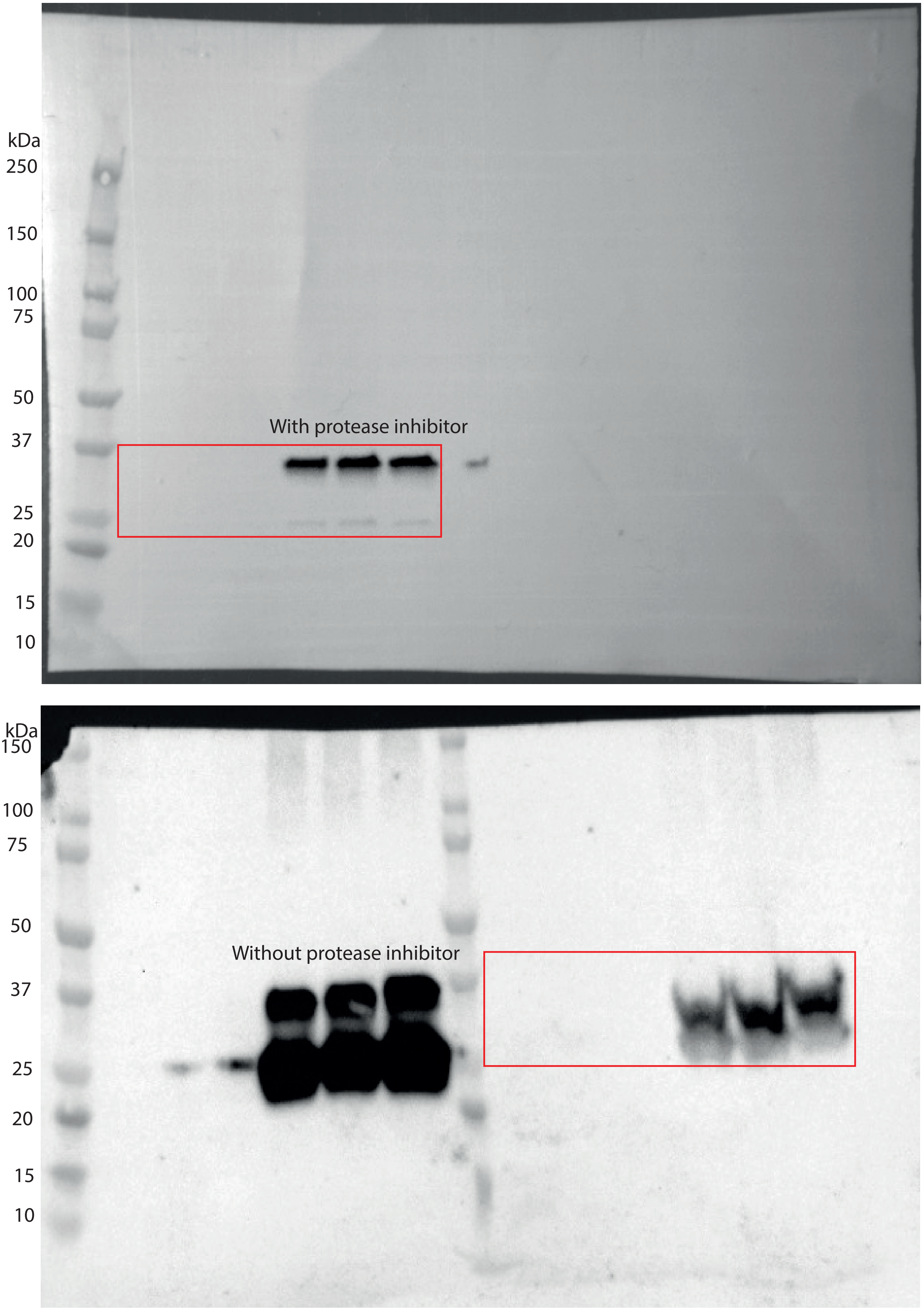

Figure S3: The orginal full-lenght EGFP on CD9-M1 cells on the upper blot and their EVs on the lower blot. Red rectangle indicate included part in figure 1.

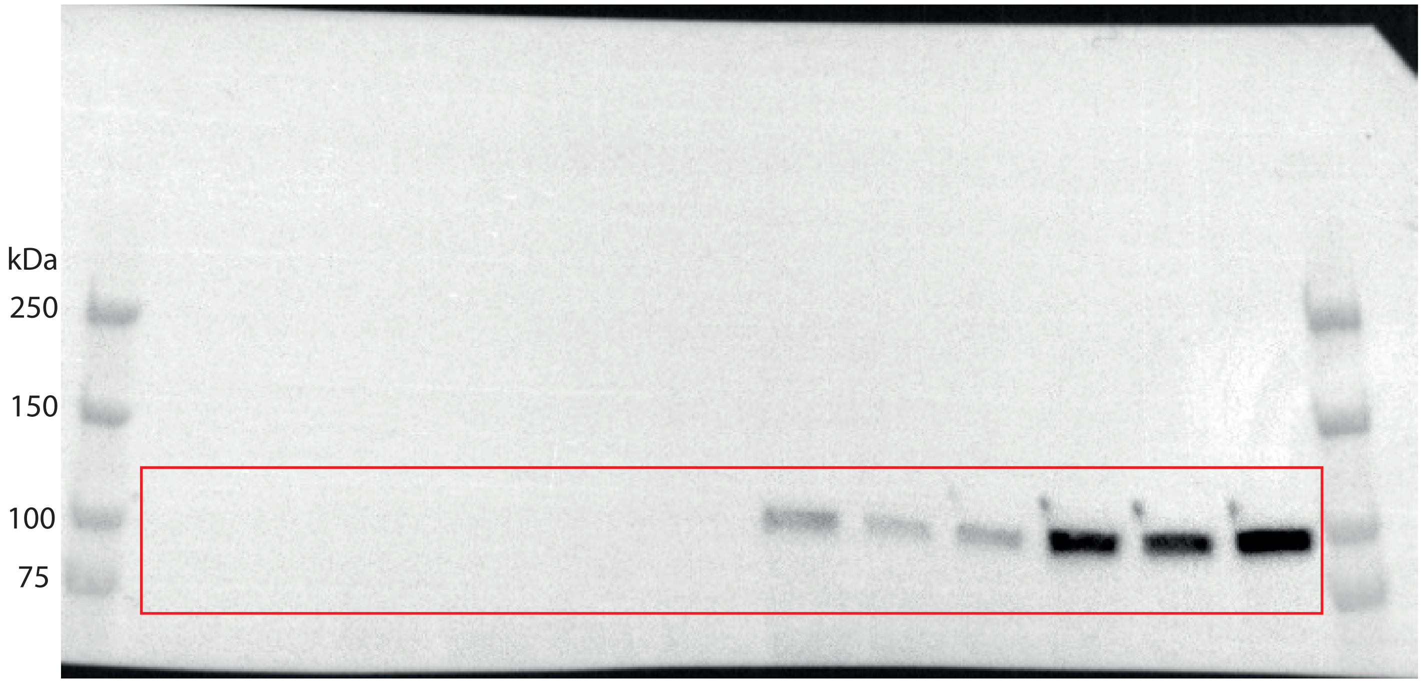

Figure S4: The orginal full-lenght ALIX blots on CD9-M1 cells and their EVs from figure 2C. Red rectangle indicate  included part in figure 2.

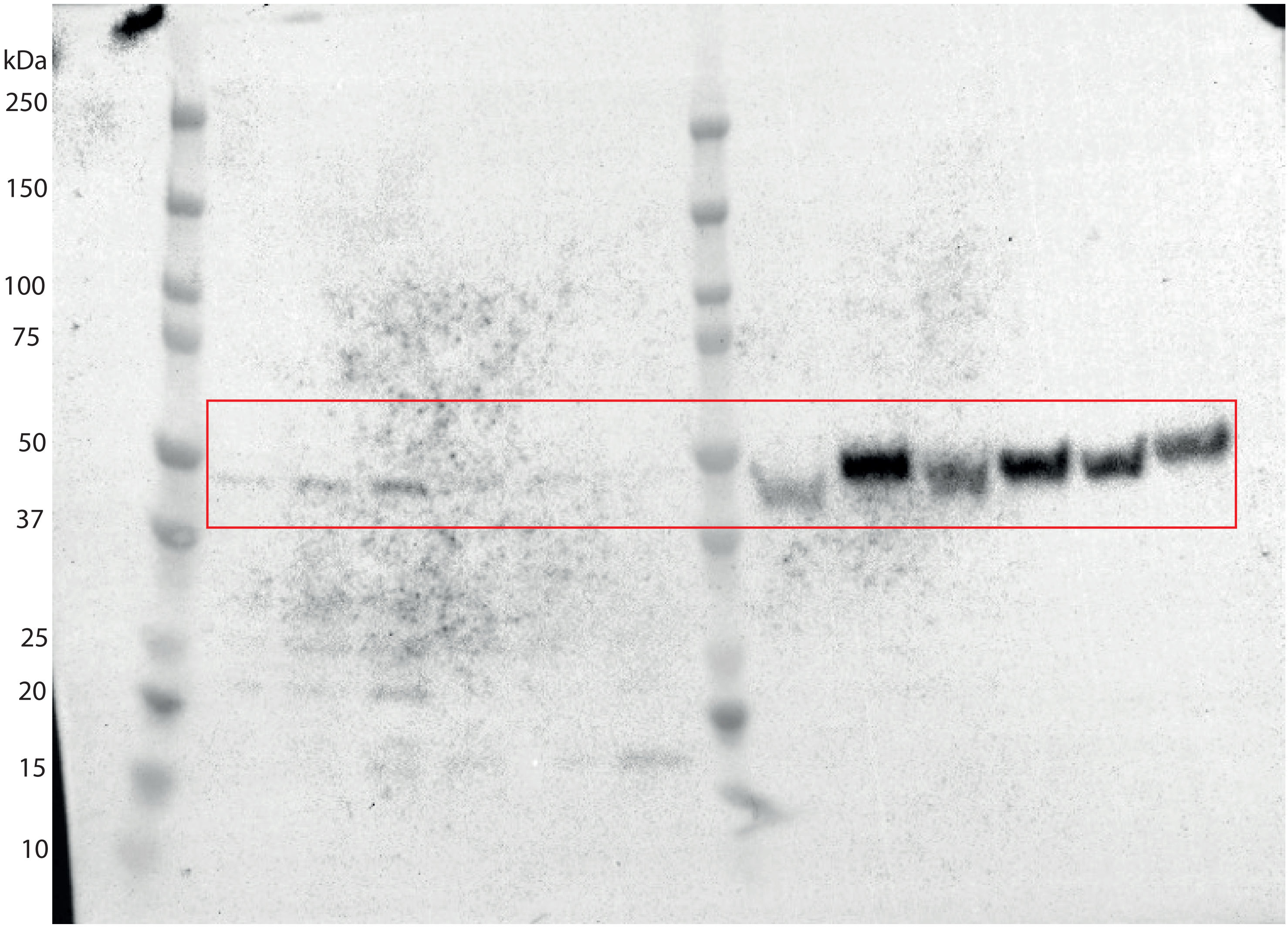

Figure S5: The orginal full-lenght Flotilin blots on CD9-M1 cells and their EVs from figure 2C. Red rectangle indicate  included part in figure 2.

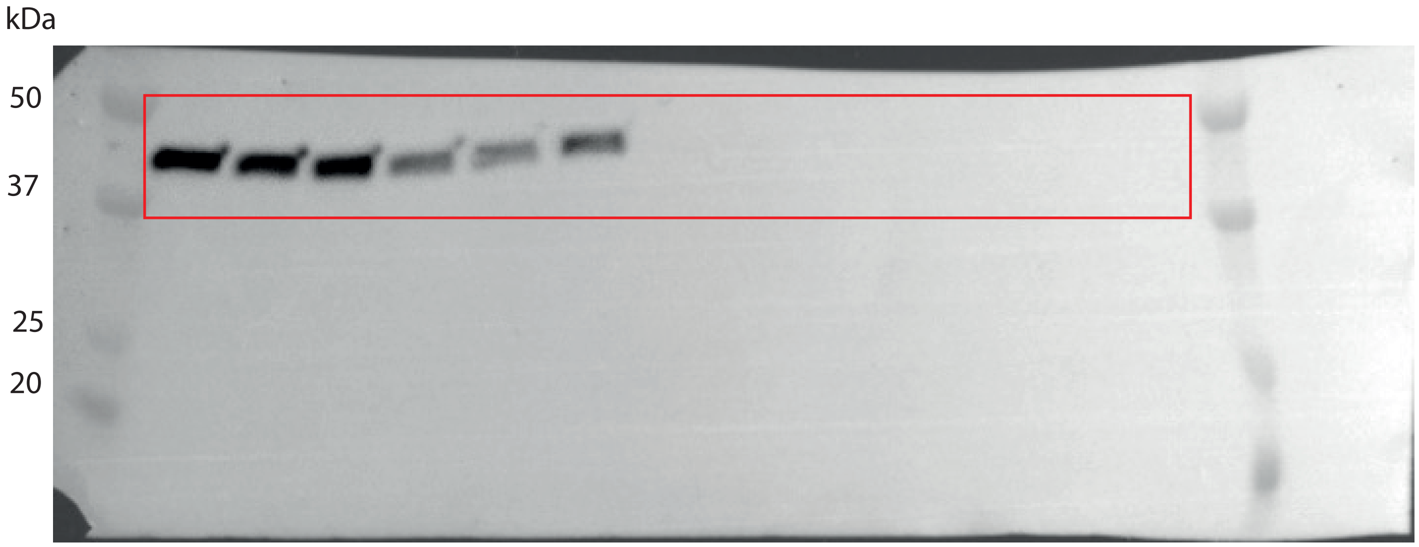

Figure S6: The orginal full-lenght Actin blots on CD9-M1 cells and their EVs from figure 2C.
Red rectangle indicate  included part in figure 2.

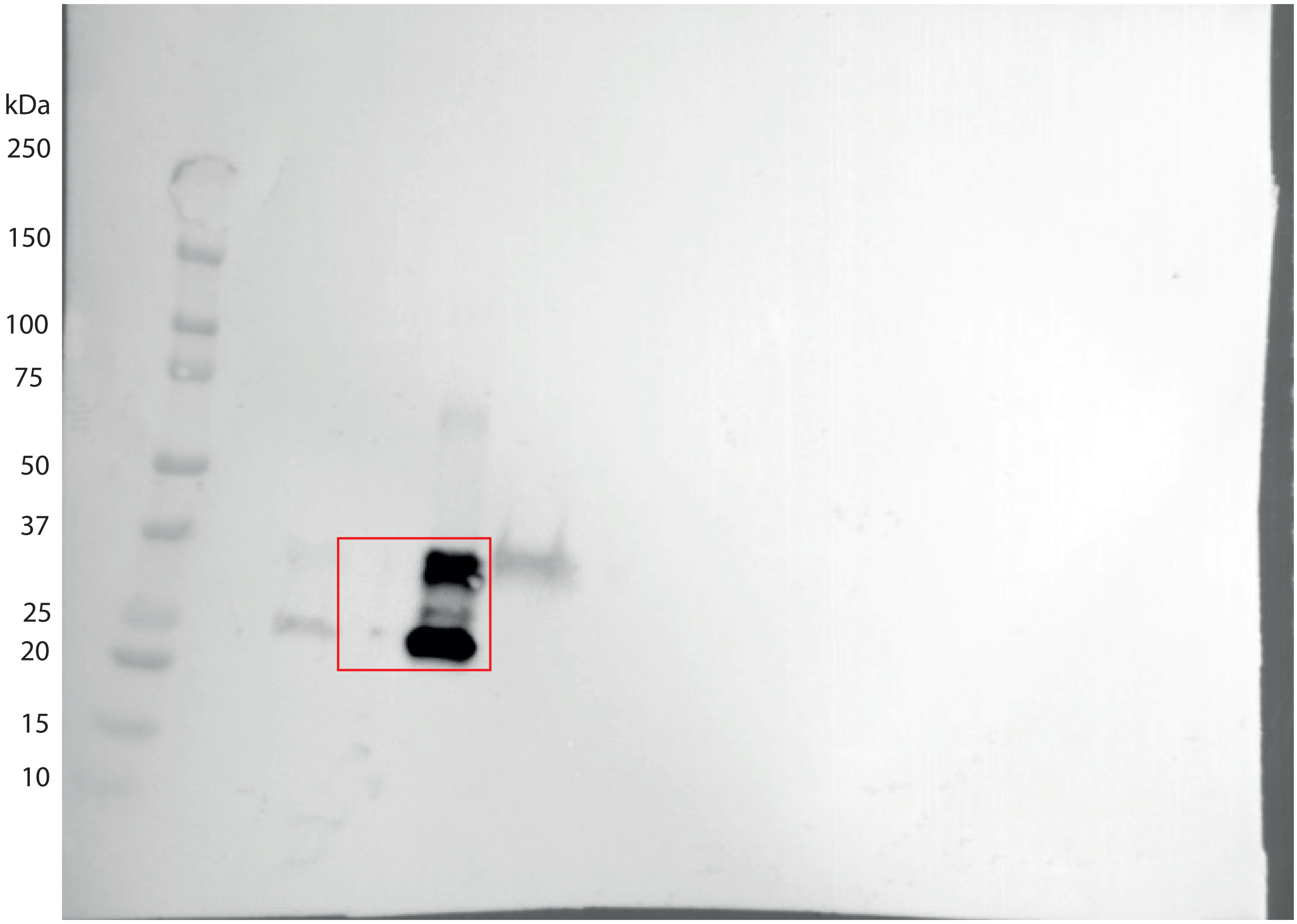

Figure S7: The orginal full-lenght EGFP blots on input and percitipated CD9-M1 medium from figure 2E. Red rectangle indicate  included part in figure 2.

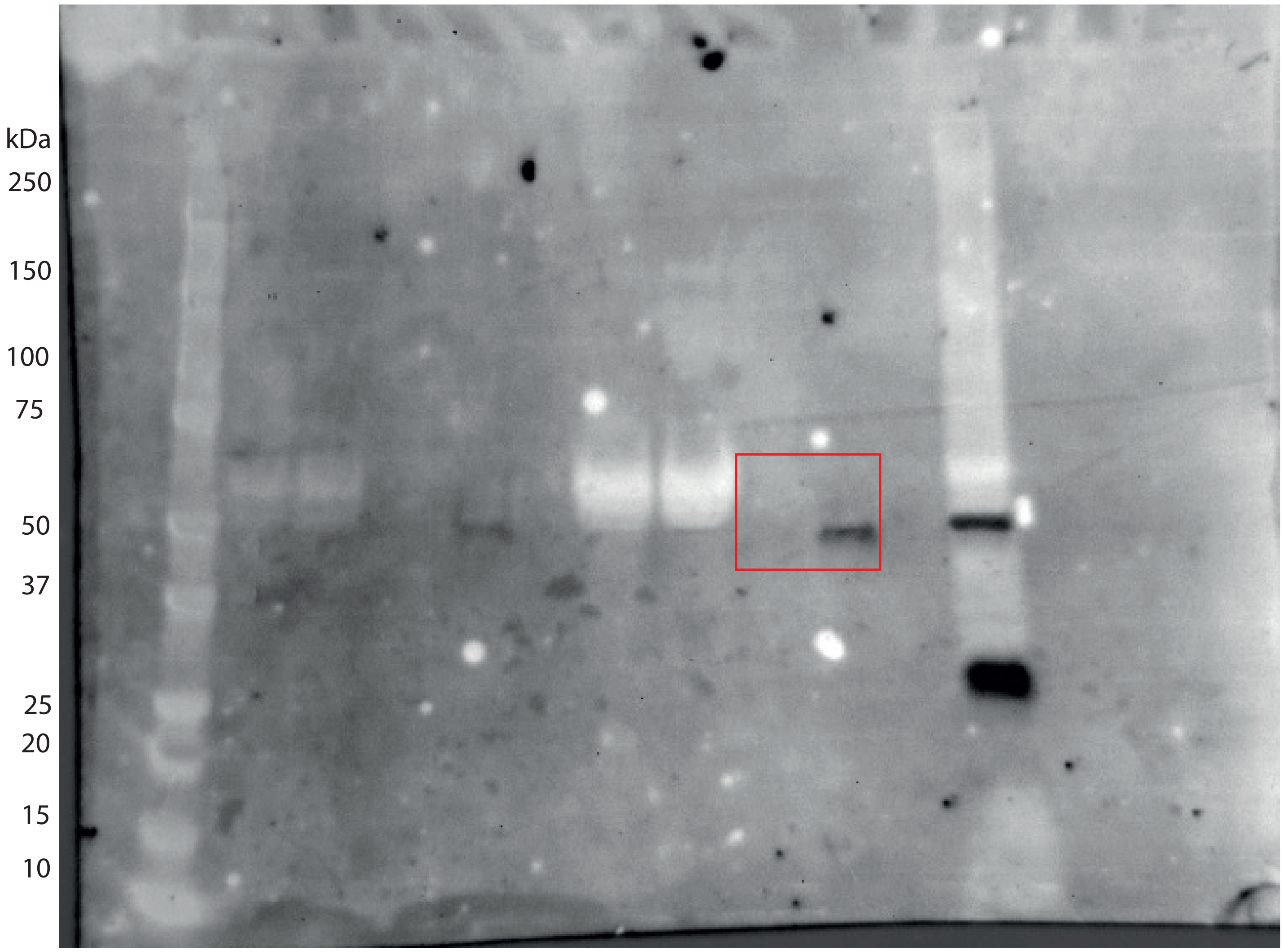

Figure S8: The orginal full-lenght TSG101 blots on input and percitipated CD9-M1 medium from figure 2E. Red rectangle indicate  included part in figure 2.

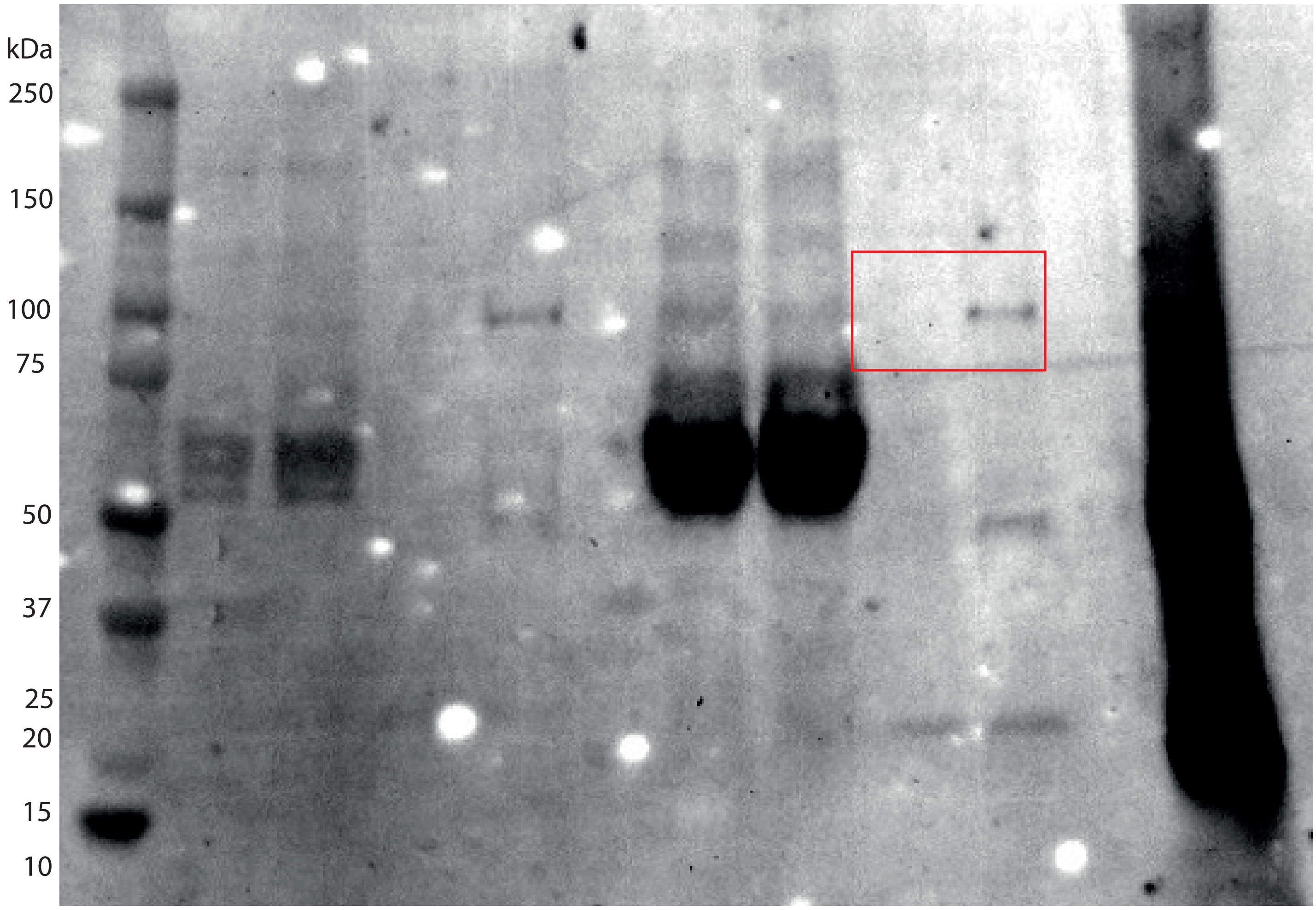

Figure 9: The orginal full-lenght ALIX blots on input and percitipated CD9-M1 medium from figure 2E. Red rectangle indicate  included part in figure 2.

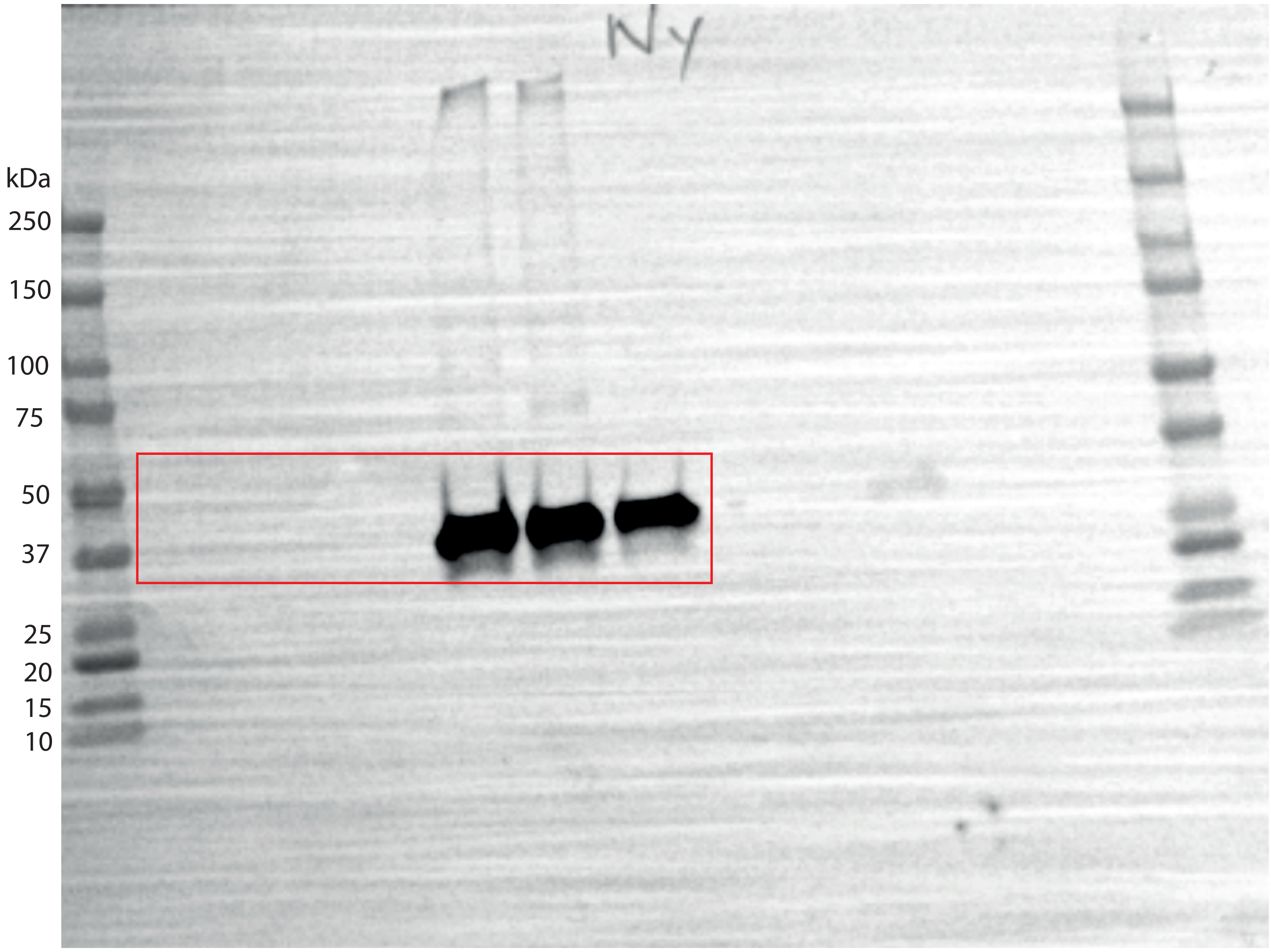

Figure S10: The orginal full-lenght EGFP blots on HEK293T cells from figure 2C. Red rectangle indicate included part in figure 2.

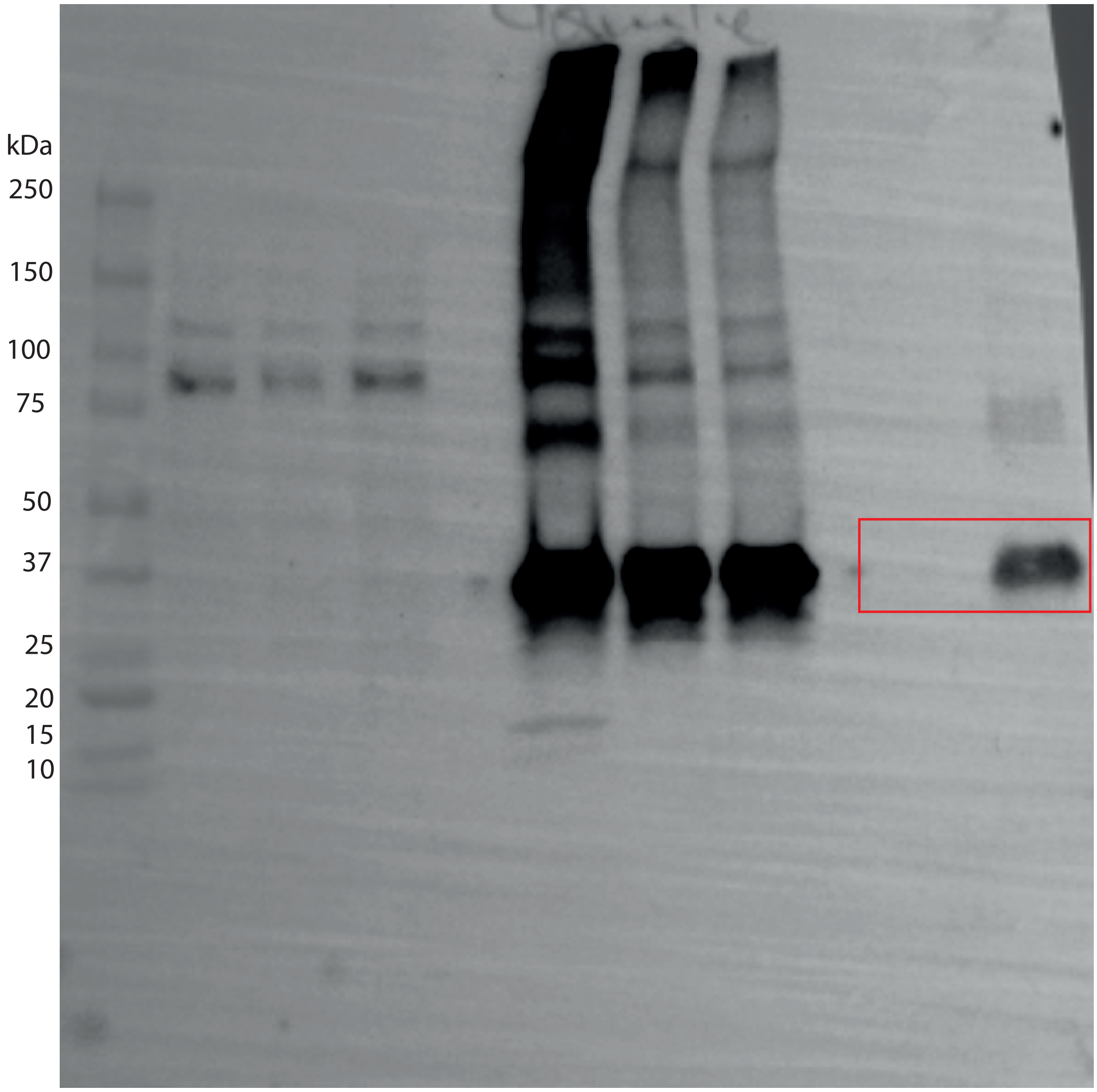

Figure S11: The orginal full-lenght EGFP blots on HEK293T medium from figure 2C. Red rectangle indicate included part in figure 3.

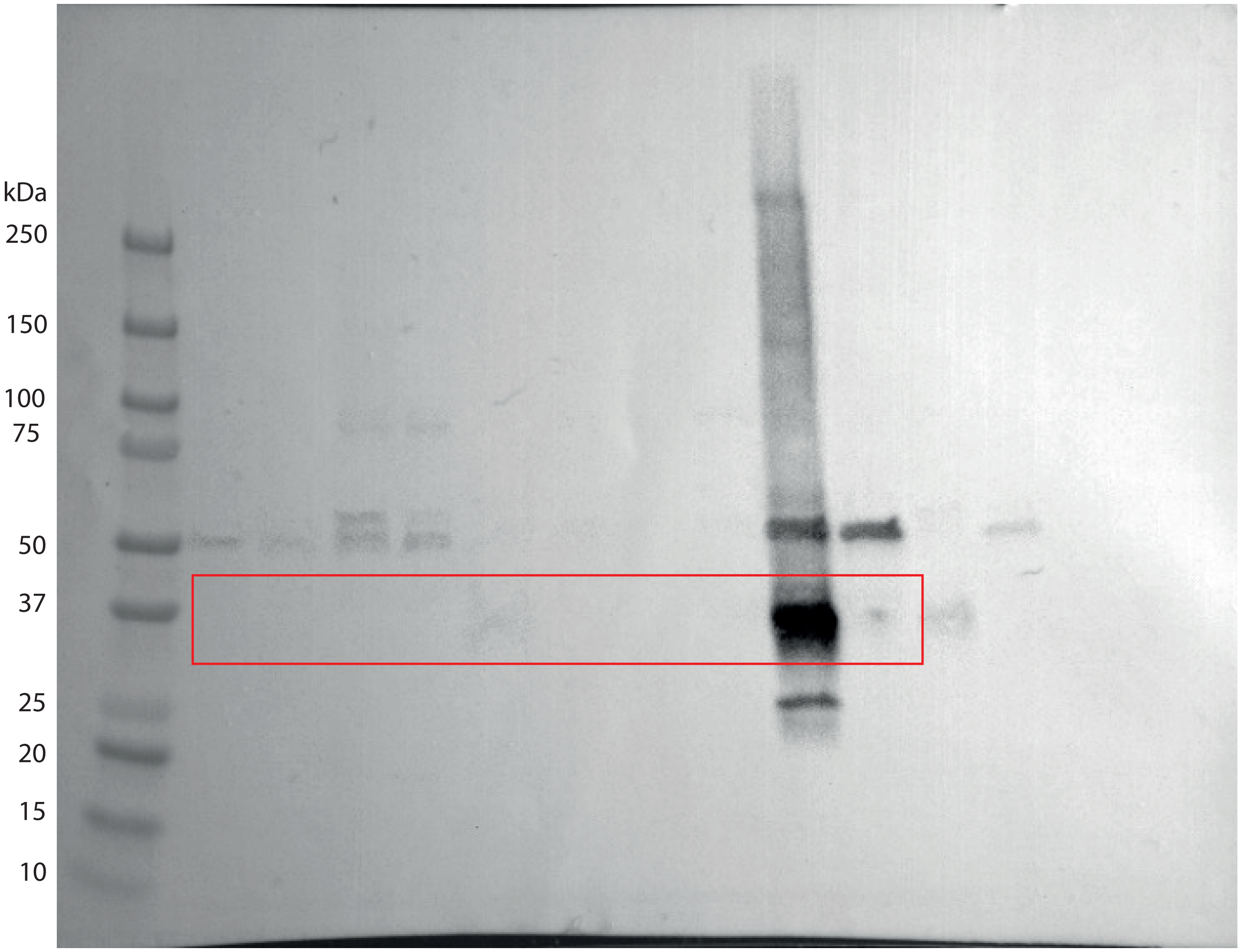

Figure S12: The orginal full-lenght GFP blots on  tissue from Kidney, Liver, Lung, Spleen and heart from figure 4A. Red rectangle indicate included part in figure 4.

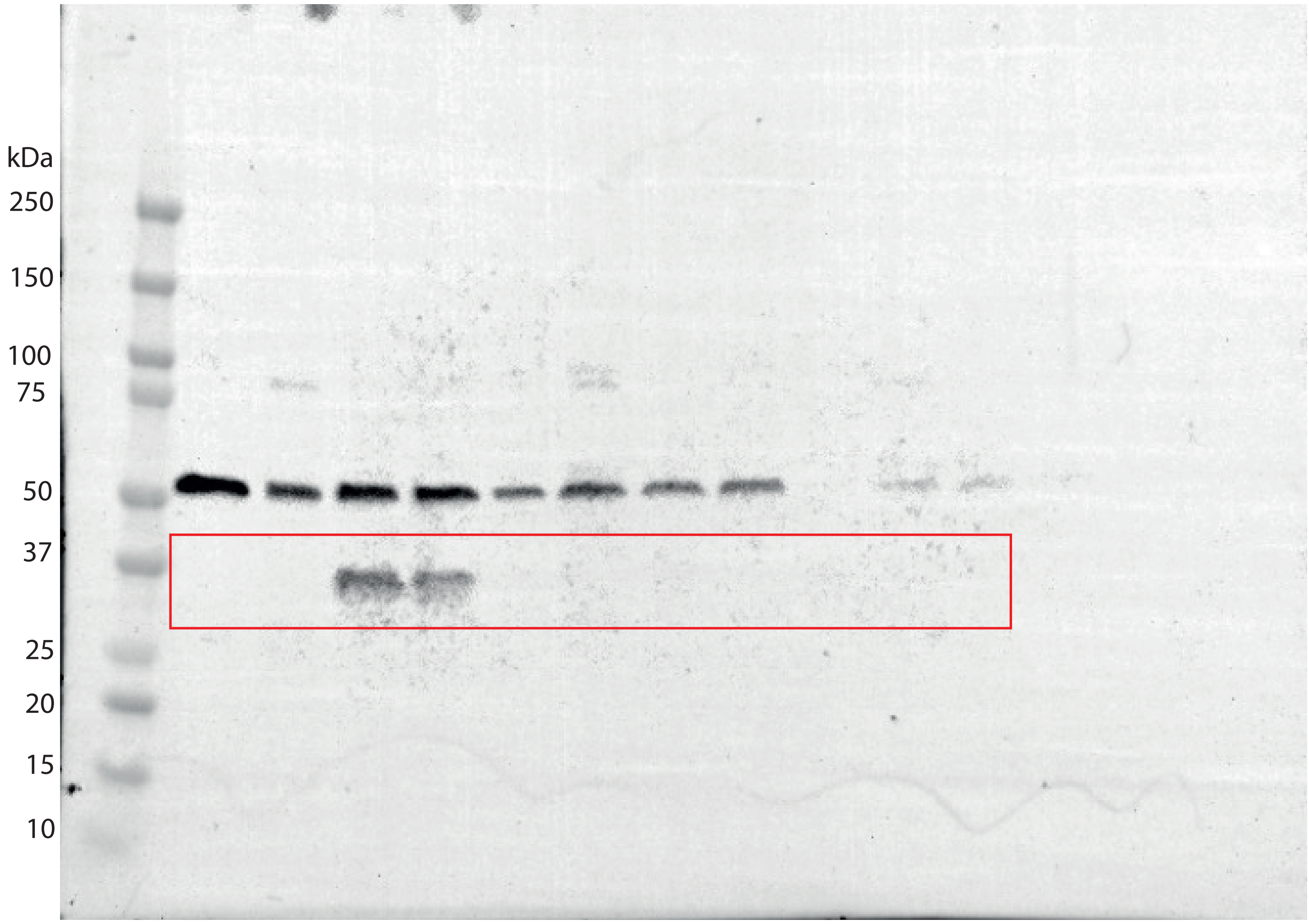

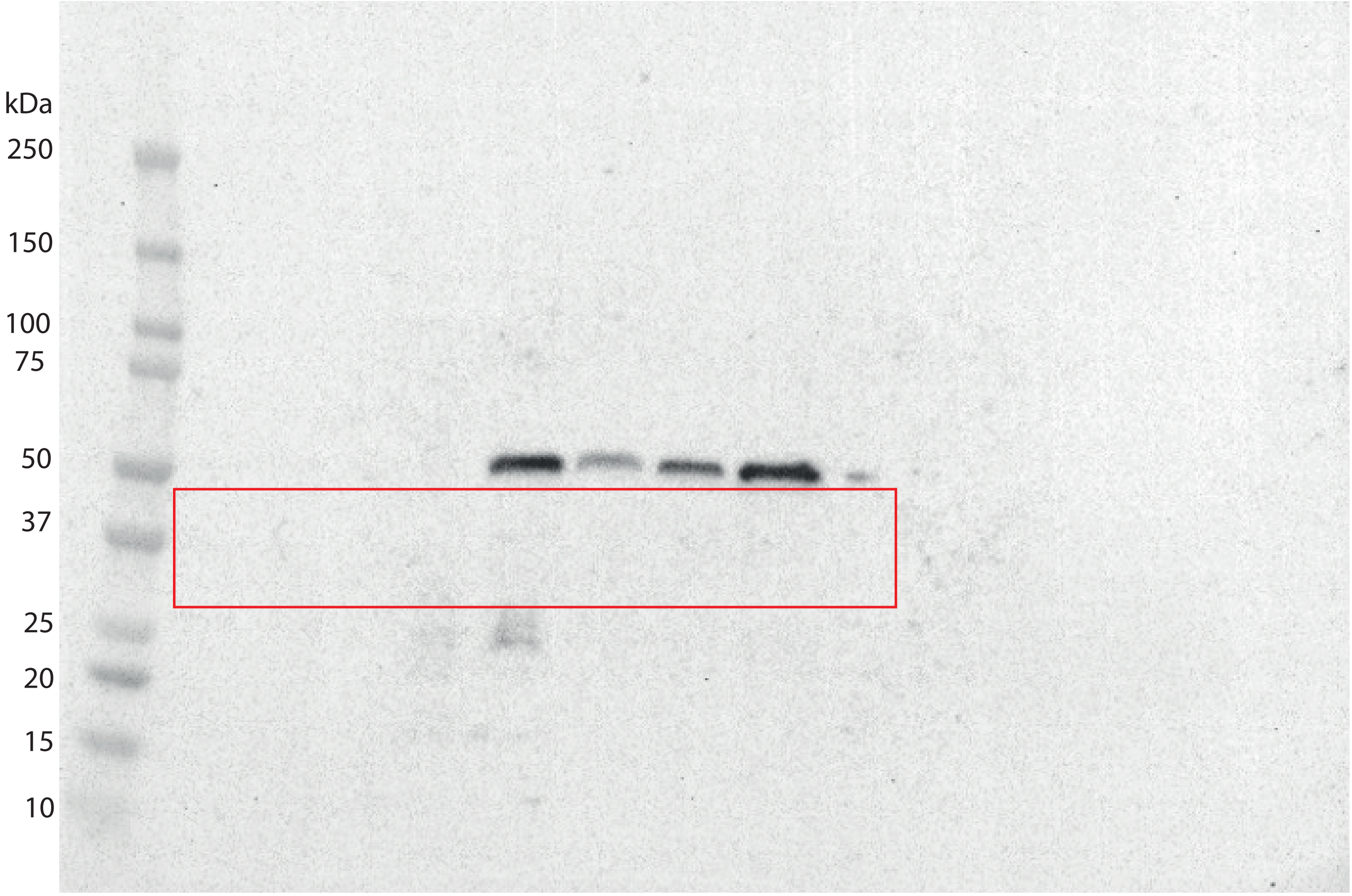

Figure S13: The orginal full-lenght GFP blots on  tissue from Kidney, Liver, Lung, Spleen and heart from figure 5A. Red rectangle indicate included part in figure 5

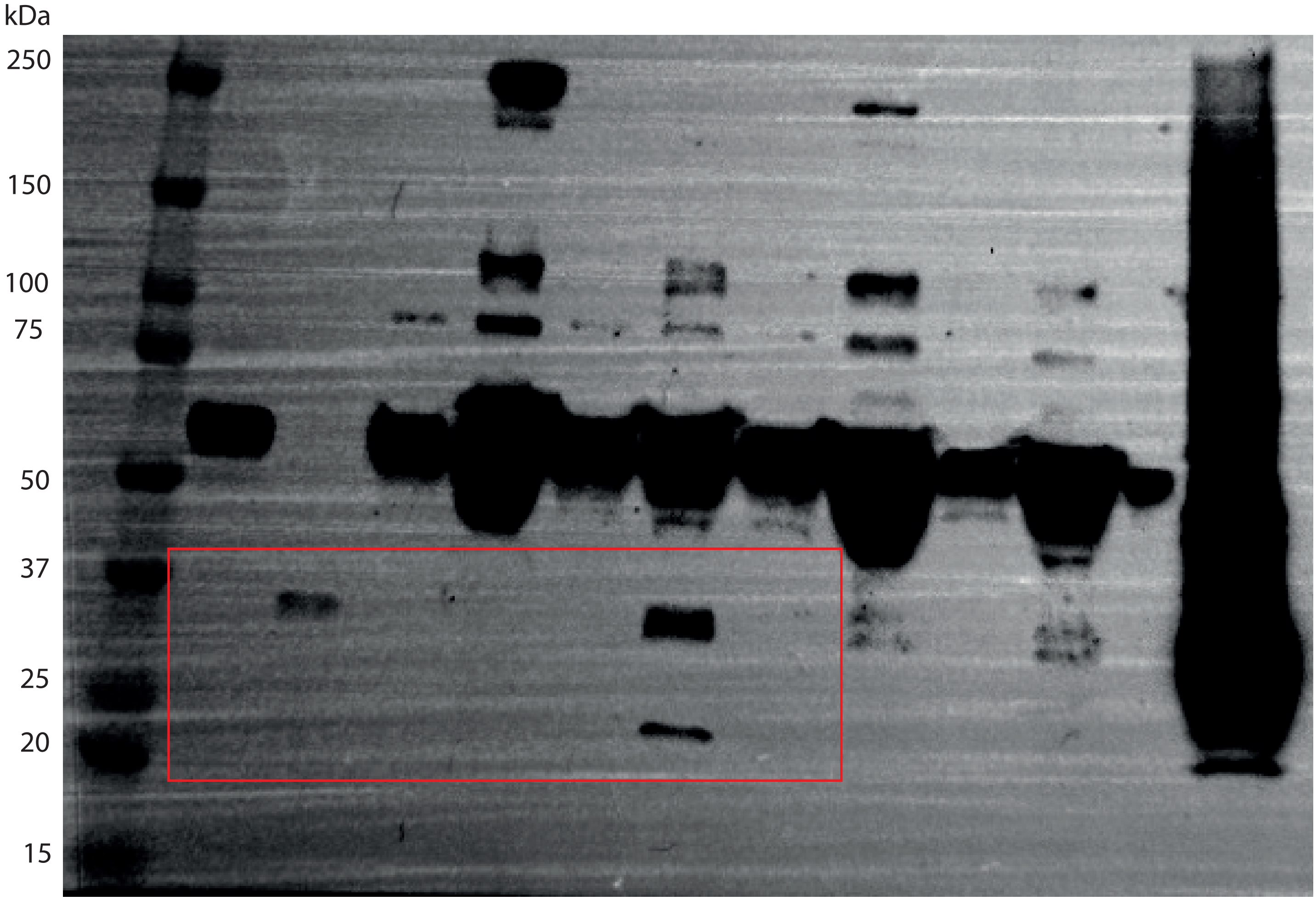

Figure S14: The orginal full-lenght GFP blot on plasma in figure 6A. Red rectangle indicate included part in figure 6.

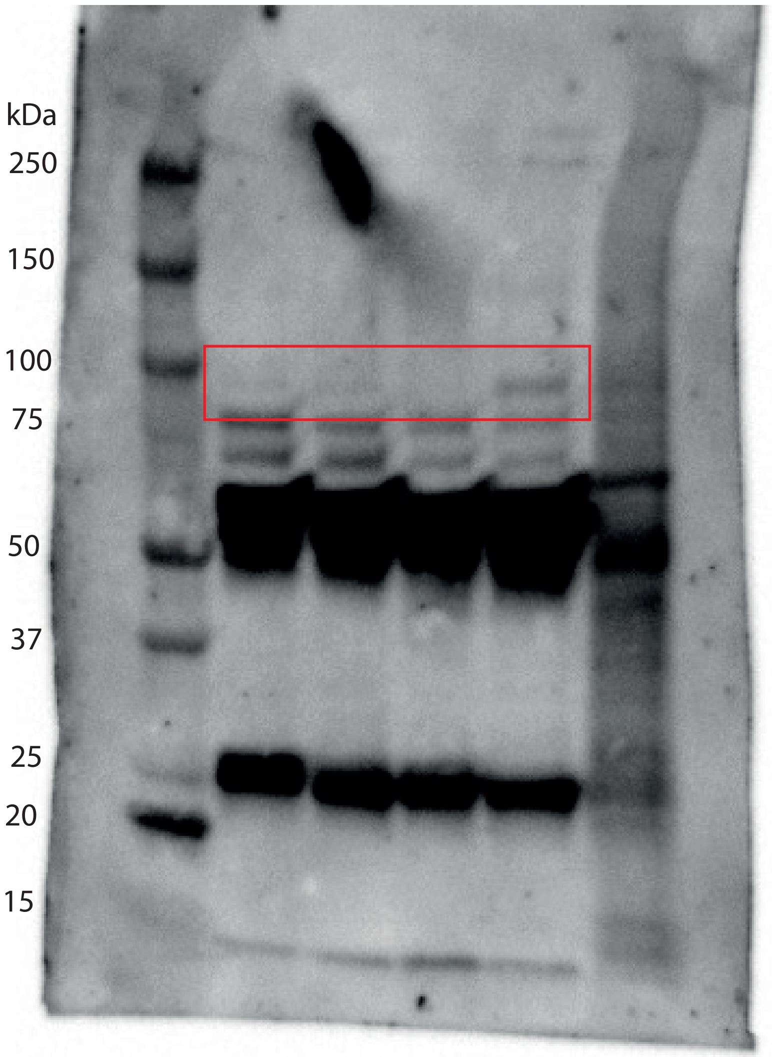

Figure S15: The orginal full-lenght ALIX blot on plasma in figure 6A. Red rectangle indicate included part in figure 6.

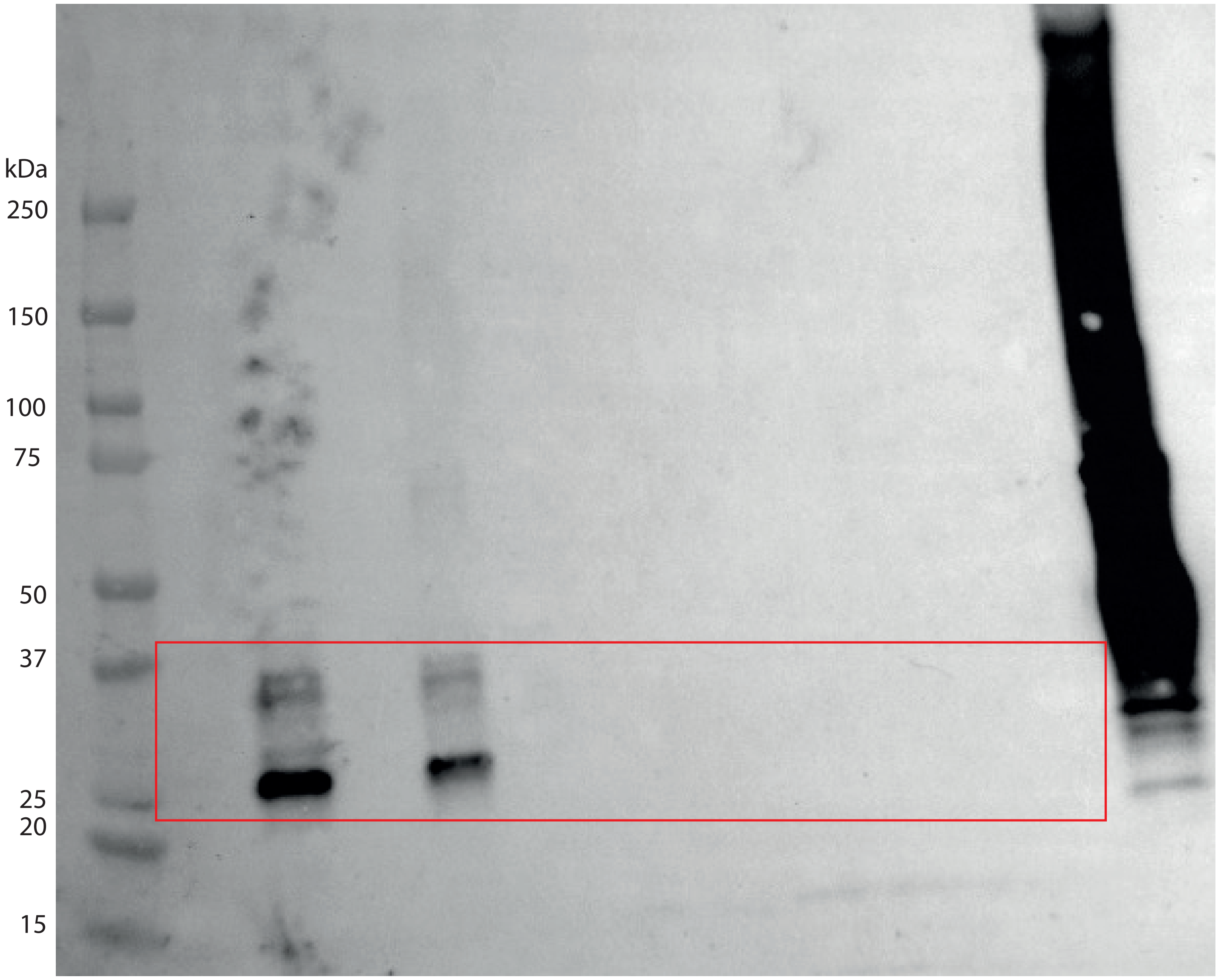

Figure S16: The orginal full-lenght GFP blots on  urine in figure 6B. Red rectangle indicate included part in figure 6.

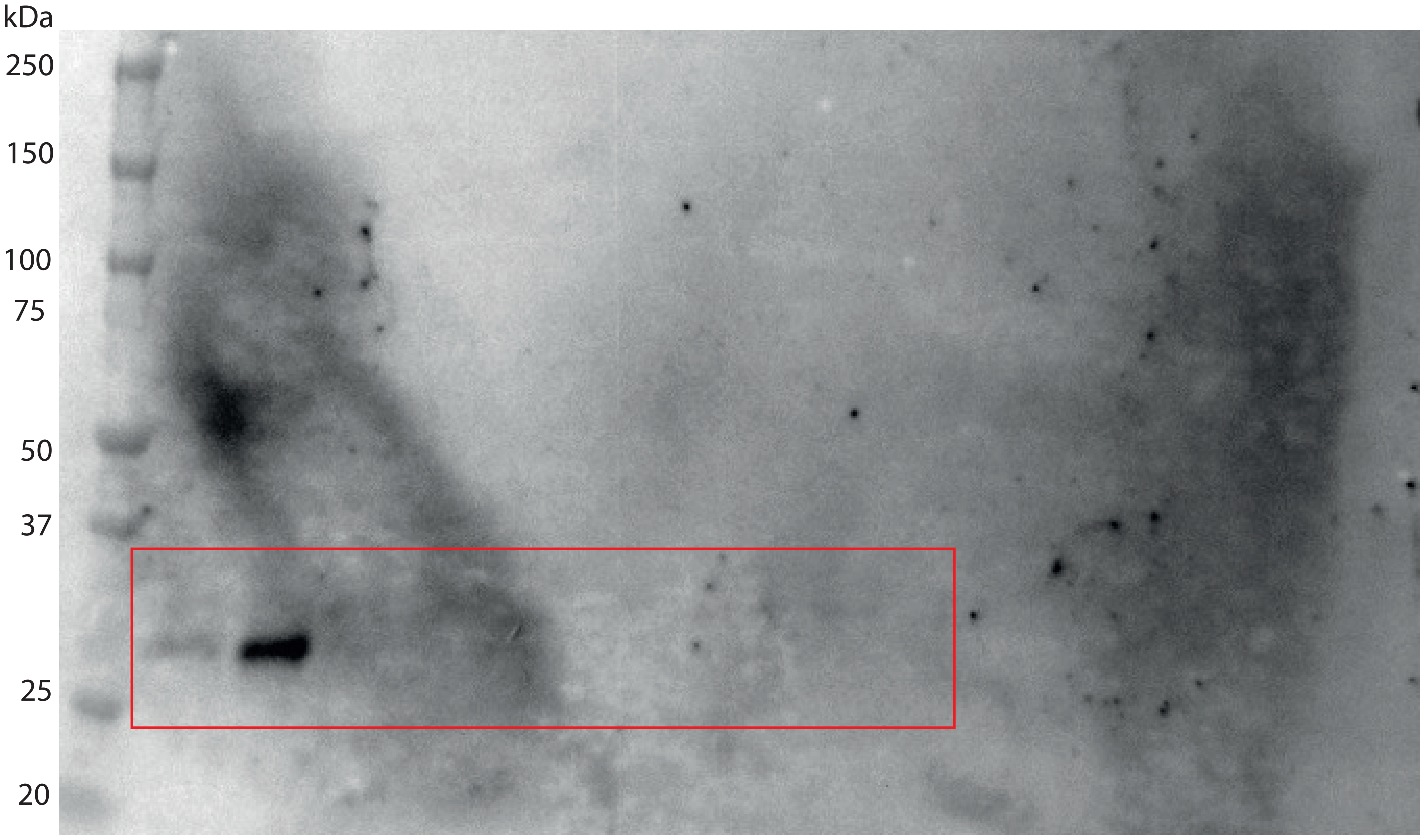

Figure S17: The orginal full-lenght CD81 blots on urine in figure 6B. Red rectangle indicate included part in figure 6.
